# Minimal cell JCVI-syn3B as a chassis to reveal the mechanisms behind *Mycoplasma* host–pathogen interactions

**DOI:** 10.1101/2022.09.19.508583

**Authors:** Daniela M. de C. Bittencourt, David M. Brown, Nacyra Assad-Garcia, Michaela R. Lynott, Lijie Sun, Luis Alberto M. Palhares de Melo, Marcelo Freire, John I. Glass

**Affiliations:** The J. Craig Venter Institute, 4120 Capricorn Lane, La Jolla, CA 92037, USA; The J. Craig Venter Institute, 9605 Medical Center Drive, Suite 150, Rockville, MD 20850, USA; Embrapa Genetic Resources and Biotechnology / National Institute of Science and Technology - Synthetic Biology, Parque Estação Biológica, PqEB, Av. W5 Norte (final), Brasília, DF, 70770-917, Brazil, Norte (final), Brasília, DF, 70770-917, Brazil

## Abstract

Mycoplasmas are atypical bacteria that are obligate parasites of animals and plants. They are known for their capacity to contaminate and parasitize mammalian cell cultures, thereby offering fortuitous models to examine the host-microbe interaction. We used a near wildtype strain of the goat pathogen *Mycoplasma mycoides* (*Mmc*) and several minimized *Mmc* strains lacking genes not essential for growth in laboratory media to investigate host-mycoplasma interactions. Stains with near minimal genomes were incapable of surviving in co-culture with mammalian HEK-293T and HeLa cells. We identified a cluster of eight non-essential genes that when added back to the minimized strains enabled mycoplasma attachment to mammalian cells. Those genes did not restore the capacity of the minimal cell to grow when co-cultured with mammalian cells indicating processes of attaching to mammalian cells and parasitism involve different genes. Furthermore, minimized *Mmc* strains were not phagocytized by human myeloid cells unlike the near wildtype *Mmc*.

## Introduction

*Mycoplasmas* were first described in 1898 by Nocard and Roux, in animals with contagious bovine pleuropneumonia (*Mycoplasma mycoides* subsp. *mycoides*). Lacking cell walls, mycoplasmas are atypical bacteria, the *Mollicutes*, named from the Latin “soft” (*mollis*) and “skin” (*cutis*) (Noccard and Roux, 1898).

Widely distributed among humans, animals, insects and plants, these microorganisms are the smallest and simplest self-replicating bacteria reported to date, with a genome size ranging from 580 kb for *Mycoplasma genitalium* (Fraser et al., 1995) to 1358 kb for *Mycoplasma penetrans* (Sasaki et al., 2002). Mycoplasma genomes evolved from bacteria similar to the model organism *Bacillus subtilis* by a process of massive gene loss. In adapting to diverse nutrient rich environments, such as the respiratory systems or urogenital tracts of mammals, mycoplasmas from several phylogenetic lineages have discarded genes for cell wall synthesis or synthesis of nucleic, amino, or fatty acids (Woese et al., 1980). In nature, because of their limited metabolic capabilities and obligate parasitic lifestyles mycoplasmas have evolved to adapt to distinctive usually highly limited niches (Benedetti et al., 2020). However that requirement for a specific host does not apply to the capacity of mycoplasmas to parasitize, i.e. become contaminates in cultured animal cells in laboratory environments.

Moreover, mycoplasma contamination poses a frequent threat to mammalian cell cultures and biological materials (Bolske, 1988; Kong et al., 2001; Steiner and McGarrity, 1983; Timenetsky et al., 2006). Experimental results can become unreliable and biologic products defective if exposed to a viable strain of mycoplasma (Nikfarjam and Farzaneh, 2012). The use of contaminated cells has the potential to invalidate results because the mycoplasma contaminants can alter almost all aspects of cell physiology. While this can lead to erroneous results or causes the loss of unique cell lines, it also offers an opportunity to examine host-parasite interactions (Nishiumi et al., 2021).

Members of the *mycoides* clade of mycoplasmas are not among the five mycoplasma species that cause ~95% of cell culture contaminations. Nor is it among the 20 mycoplasma species reported as cell culture contaminants (Tang et al., 2000). Still these mycoplasmas are important veterinary pathogens that cause diseases that are often chronic in nature (Rottem, 2003). For instance, the pathogenic members of the *mycoides* clade can cause serious livestock diseases (7). *Mycoplasma. mycoides* subsp. *mycoides* is the causative agent of contagious bovine pleuropneumonia (1) and *M. mycoides* subsp. *capri* (*Mmc*) causes the “MAKePS” syndrome, characterized by mastitis, arthritis, keratoconjunctivitis, pneumonia and septicemia in goats (Thiaucourt and Bolske, 1996). Both strains can result in outbreaks with high mortality (Hernandez et al., 2006; Tambi et al., 2006). Because of their importance as veterinary pathogens, members of the *mycoides* clade of mycoplasmas have long been investigated to determine their virulence mechanisms and how to prevent and treat infections. Still many of the molecular mechanisms of mycoplasma infection are unknown (Di Teodoro et al., 2020; Di Teodoro et al., 2018). Here we used a system comprised of mycoplasmas and cultured mammalian cells to obtain basic information about these interactions.

The lack of information about the molecular mechanisms of pathogenicity of *M. mycoides* makes design of safe and efficient vaccines more difficult. Most of the commercial mycoplasma vaccines were developed using empirical approaches and often cause side effects and have limited efficacy and duration of immunity (Nicholas et al., 2009).

Bacteria with synthetic genomes, representing native pathogens or their fully or partially minimized counterparts, can be used to define genes responsible for specific microbe-host interactions. Mycoplasmas, as the self-replicating bacteria with the smallest genomes known have been proposed as possible guides for the design of synthetic organisms (Morowitz, 1984). Towards that goal, in 2010 a 1.1 Mb synthetic genome based on the genome of *Mmc* GM12 was used to create the strain JCVI-syn1.0 (Gibson et al., 2010). That strain contains almost all the genes encountered in the naturally occurring organism. More recently, the JCVI constructed a minimized *Mmc* cell, JCVI-syn3.0, that was built using several cycles of genome design that eliminated more than half of the genes in JCVI-syn1.0, resulting in a bacterium with a 531 kb genome encoding 473 genes (Hutchison et al., 2016). The JCVI-syn3.0 has the smallest genome of any known organism that can be grown in axenic culture. We have also constructed variants of minimal cell JCVI-syn3.0 that have a small number of added non-essential genes that result in a cell with properties more like wild type *Mmc* (Breuer et al., 2019; Hossain et al., 2021; Pelletier et al., 2021).

In 2021 we reported that a minimized *Mmc* strains JCVI-syn3.0, JCVI-syn3A and JCVI-syn3B were unable to survive in co-culture with host cells such as HeLa (Nishiumi *et al*., 2021). The JCVI-syn3A (Breuer *et al*., 2019) and JCVI-syn3B strains differ from JCVI-syn3.0 by the presence of 19 additional non-essential genes that result in a more easily manipulated cell. JCVI-syn3B additionally includes a dual *loxP* landing pad that enables easy Cre recombinase mediated insertion of genes. None of the added 19 genes are expected to alter the pathogenicity of JCVI-syn3A or JCVI-syn3B. Rather they make the strains more robust for laboratory operations and give the cells a more normal *Mmc* phenotype. Furthermore, addition of a surface antigen gene called *mba* from the mycoplasma species *Ureaplasma parvum* serovar 3, restored the capacity of JCVI-syn3B to adhere to HeLa cells. We also showed that a JCVI-syn3B construct expressing *U. parvum mba* and a *U. parvum* gene proposed to be a virulence factor blocked endoplasmic reticulum stress-induced cell death, caused the formation of vacuoles in the HeLa cells and enabled proliferation of the mutant strain in co-culture with the cell host (Nishiumi *et al*., 2021).

The present study uses assorted analytical methods to confirm that the minimal cell JCVI-syn3.0 and its similarly minimized derivatives cannot survive in co-culture with mammalian cells. To do this we utilized assays that detected viable mycoplasmas after one to two weeks of co-incubation with host mammalian cells or based on PCR detection of mycoplasmas after such incubations. Note that detection of living mycoplasmas or mycoplasma DNA does necessarily mean the parasites were capable of proliferating under the experimental conditions, only that they were still present and alive. The difference between survival and proliferation is an important distinction. Our experiments describe mycoplasma cells that were incubated with mammalian cells either as not surviving or having a survival or proliferation phenotype. The survival phenotype refers to bacteria that lived more than 10 days of incubation with mammalian cells, but did not proliferate during that incubation to any measurable amount. Proliferation indicates the bacterial cells are actively parasitizing their mammalian cell hosts.

Additionally, comparison of the genomes of all the strains tested identified a cluster of eight candidate genes that might be correlated with their survival when co-cultured with mammalian cells. To effectively parasitize host mammalian cells, the *Mmc* strains must be able to adhere to and proliferate while attached to their host cells. To better understand the genetic factors and mechanism behind the capacity of *Mmc* to survive in co-culture with mammalian cells and, perhaps, also pathogenicity, we inserted these eight genes back into the dual *loxP* landing pad in JCVI-syn3B strain (Nishiumi *et al*., 2021; Sauer, 1987) (individually and the complete cluster) and evaluated their contribution for *Mmc* adherence and proliferation with the mammalian cell hosts. Our experiments determined that those eight non-essential genes restored the capacity of the minimized *Mmc* strains to attach to mammalian cells but not the proliferation phenotype.

Additionally, we hypothesized that because most of the proteins associated with the *Mmc* membrane were not present on the minimal cell, the minimal cell might not be detected by all or some of the elements of mammalian immune systems. We tested this hypothesis by incubating both the near wild type *Mmc*-like synthetic strain JCVI-syn1.0 and various minimized versions of *Mmc* with neutrophil-like differentiated human promyelocytic leukemia cells (dHL-60). These dHL-60 cells readily phagocytized the wild type organisms but not the minimized cell, JCVI-syn3A. We also demonstrated the JCVI-syn3B strain expressing the eight *Mmc* non-essential genes that enabled the mutant to attach to mammalian cells induced a higher phagocytic activity. These studies demonstrating the low immunogenicity of the minimal *Mmc* suggest possible uses for the organism as a vehicle for delivering therapeutic proteins or small molecules.

Collectively, these studies indicate that our minimized *Mmc* strains can be used to investigate the mechanisms of mycoplasma pathogenesis and host-microbial interactions. Furthermore, because JCVI minimal bacterial cells did not survive in co-culture with mammalian cells and were not phagocytized by neutrophils, we speculate these minimized bacterial might be used as chassis to express and investigate the pathogenic potential of bacterial genes encoded by genetically intractable bacterial pathogens.

## Results

### Mammalian cell culture media and conditioned mammalian cell culture growth media do not support synthetic Mmc growth

Cultured mammalian cells consume nutrients from their growth media and secrete a number of different natural products and proteins into that media. We tested whether mammalian cells and/or their secreted products were required for our various *Mmc* derived strains to survive in cultures of either HeLa cells or HEK-293T cells. For that, fresh and conditioned Dulbecco’s modified Eagle’s medium (DMEM) + 10% fetal bovine serum (FBS) media were infected with each strain. After 7 days of infection, no media or conditioned media samples that were inoculated with JCVI-syn1.0 or minimized *Mmc* strains showed any signs of bacterial metabolism. Thus, the *Mmc* strains were unable to survive or proliferate in either fresh or conditioned cell culture media.

### Detection of synthetic Mmc cell in co-culture with mammalian cells

To evaluate if our mycoplasma synthetic strains could survive and proliferate in co-culture with mammalian cells we did an infectivity assay. The JCVI-syn1.0 is a non-minimized synthetic strain based on the genome sequence of *Mmc*; although, some potential virulence related genes are not present in this organism (Gibson *et al*., 2010). On the other hand, JCVI-syn2.0 is a minimized synthetic strain with a genome size of 576 kb and 516 genes, 47 more than the minimal cell JCVI-syn3.0 (Hutchison *et al*., 2016). We thought it was highly likely that the genes responsible for infectivity and pathogenesis have been removed from JCVI-syn2.0. We speculated that the infectivity and pathogenesis genes may also be involved in the capacity of *Mmc* to parasitize mammalian cells in tissue culture.

Once we confirmed JCVI-syn1.0 and our minimized *Mmc* strains could not grow in mammalian medium, we demonstrated JCVI-syn1.0 was capable of proliferating in cultures of HEK-293T cells. Contamination was evaluated at select time points after mixing the prokaryotic and eukaryotic cells and after several passages of the mammalian cells. We determined JCVI-syn1.0 survived in mammalian cell cultures, it also proliferated. We titrated mycoplasmas 14 and 15 days after they mixed with mammalian cells (5^th^ passage of culture medium) and found a 10-100-fold increase in Color Changing Units (CCUs – a method commonly used to titer mycoplasmas after the additional 24 h incubation) (**Figure S2**).

JCVI-syn2.0 was evaluated by the same method used to show that JCVI-syn1.0 grew in co-culture with HEK-293T cells (**Table 1**). We detected JCVI-syn2.0 for 5 days post infection and before the 2^nd^ passage of HEK-293T cells culture medium but not at 7 days. Results from this infectivity assay formed the basis for determining the threshold of detecting whether each mycoplasma strain was able to survive in mammalian culture. If mycoplasma contamination was present after 7 days post infection and after two passages, strains were designated as possessing a survival phenotype. We could not detect mycoplasma contamination using an inverted microscope at 30x magnification (**Figure S3**).

**Table 1.** CCU assay of Synthetic *Mmc* persistence in HEK-293T cells. Actively growing cultures of JCVI-syn 1.0 or JCVI-syn 2.0 were added to monolayer cultures of HEK-293T cells. At selected times for 15 days (d) samples of supernatants were removed and a CCU assay was performed. Mycoplasma was detected (+) when a color change was identified. Passage (P#) number is given in parentheses. Pictures of the CCU assay can be seen in supplementary Figure S2.

| Strain | 1 d<br>post<br>infecti<br>on<br>(P<sub>0</sub>) | 3 d<br>post<br>infecti<br>on (P<sub>0</sub>) | 5 d<br>post<br>infecti<br>on (P<sub>1</sub>) | 7 d<br>post<br>infecti<br>on (P<sub>2</sub>) | 11 d<br>post<br>infecti<br>on (P<sub>3</sub>) | 13 d<br>post<br>infecti<br>on (P<sub>4</sub>) | 14 d<br>post<br>infecti<br>on (P<sub>5</sub>) | 15 d<br>post<br>infecti<br>on (P<sub>5</sub>) |
| --- | --- | --- | --- | --- | --- | --- | --- | --- |
| JCVI-syn<br>1.0 | + | + | + | + | + | + | + | + |
| JCVI-syn<br>2.0 | + | + | + | - | - | - | - | - |
| No<br>infection | - | - | - | - | - | - | - | - |

### Using minimized Mmc strains to identify candidate genes necessary for infection

JCVI-syn2.0 lacks about half of the genes encoded by JCVI-syn1.0 (Hutchison *et al*., 2016). To evaluate which genes or clusters of genes are important for infectivity of mycoplasmas in mammalian cell culture, we used our library of partially minimized strains in hopes of determining or narrowing down which genes in JCVI-syn1.0 genome enabled cells to survive and proliferate in co-culture with mammalian cells (**Table 2**).

**Table 2.** Infectivity assay results for synthetic *Mmc* strains. Actively growing cultures of synthetic *Mmc* were used to inoculate either a culture of HEK-293T cells; a culture of HeLa cells; or fresh DMEM media supplemented with 10% FBS; filter-sterilized spent DMEM media supplemented with 10% FBS after being used as growth medium for 2 days with HEK-293T or HeLa cells. Seven days post infection a sample of supernatant or medium was removed and a CCU assay was performed. Mycoplasma was detected (+) when a color change was identified.

| Strain | HEK-293T | HeLa | Unspent DMEM media | HEK-293T spent media | HeLa spent media |
| --- | --- | --- | --- | --- | --- |
| JCVI-syn 1.0 | + | + | - | - | - |
| JCVI-syn 2.0 | - | - | - | - | - |
| JCVI-syn 3.0 | - | - | - | - | - |
| syn 1.0F | - | - | - | - | - |
| RDG1.0-1 | + | n/a | n/a | n/a | n/a |
| RDG1.0-2 | - | n/a | n/a | n/a | n/a |
| RDG1.0-3 | + | n/a | n/a | n/a | n/a |
| RDG1.0-4 | + | n/a | n/a | n/a | n/a |
| RDG1.0-5 | + | n/a | n/a | n/a | n/a |
| RDG1.0-6 | + | n/a | n/a | n/a | n/a |
| RDG1.0-7 | + | n/a | n/a | n/a | n/a |
| RDG1.0-8 | + | n/a | n/a | n/a | n/a |

n/a stands for not attempted

First we used the HEK-293T cell infectivity assay on a set deletion mutants made in the process of creating JCVI-syn3.0 called the Reduced Genome Design (RGD1.0) strains (RGD1.0-X refers to the set of 8 strains labeled RGD1.0-1, RGD1.0-2, etc) (Hutchison *et al*., 2016). Each of the eight RGD1.0 strains has only 1/8^th^ of its genome minimized. For example, RGD1.0-3 has 7/8^ths^ of its genome from JCVI-syn1.0 and the 3^rd^ segment of the genome from JCVI syn 2.0 (**Figure S4**). This 8-strain series was used to systematically test each 1/8^th^ segment for the survival phenotype. Testing each of the RGD1.0 series showed only RGD1.0-2 lost the capacity to survive and grow in co-culture with mammalian cells, meaning the deleted genes from segment 2 resulted in a loss of the survival and proliferation phenotypes. Then, we analyzed another minimized strain that also lost the capacity to proliferate in mammalian cell cultures, designated as syn1.0F. The difference between syn1.0F and JCVI-syn1.0 is syn1.0F lacked a cluster of 8 segment 2 genes. The genes are MMSYN1_0179, MMSYN1_0180, MMSYN1_0181, MMSYN1_0182, MMSYN1_0183, MMSYN1_0184, MMSYN1_0185 and MMSYN1_0186. Their annotations are listed in Table 3. Genes 0179 through 0184 encode elements of a likely ABC sugar transporter, perhaps of maltose or some other disaccharide (based on current GenBank annotations, and annotations of orthologous genes in related mycoplasma species). All strains evaluated lacking these 8 genes, including JCVI-syn3.0, resulted in a loss of the survival phenotype (**Table 2**).

### Validation of CCU assay with PCR

Although the CCU assay has long been a standard assay for detecting mycoplasmas in cell culture and requires only one living mycoplasma cell in order to obtain a positive result, it was possible that contamination from another source led to positive CCU assays. To validate that the CCU assay was detecting our full and minimized genome strains of *Mmc*, a multiplex PCR assay was used to detect complete genomes of JCVI-syn1.0. Multiplex PCR amplification on 8 sites distributed across the 1.1 Mb *Mmc* genome was based off a previously reported multiplex PCR protocol and primers for detecting only JCVI-syn1.0 and its derivatives (Lartigue et al., 2009). Each primer set (**Table S2**) amplifies a segment at least 100 kb from any other set. The PCR target are distributed around the entire 1.1 Mb genome. Syn1.0F was derived from JCVI-syn1.0 so, if present, as the primers were designed, each syn1.0F segment would be detected.

We assayed samples for *Mmc* sequences using multiplex primers just prior to passage (i.e. supernatant sampling) on the 3^rd^ day post infection; and after the 2^nd^ such passage on the 7^th^ day post infection. From previous results we knew that cells that cannot grow in co-culture with mammalian cells on the 7^th^ day post inoculation will not result in a positive CCU assay (Nishiumi *et al*., 2021), but there are still enough cells present in the inoculum to result in a positive CCU assay on the 3^rd^ day post infection. We observed JCVI-syn1.0 was detected by PCR 7 days post infection, while Syn1.0F was not detected. These results confirmed the results from our infectivity assay evaluated via CCU (**Figure 1**).

**Figure 1.**
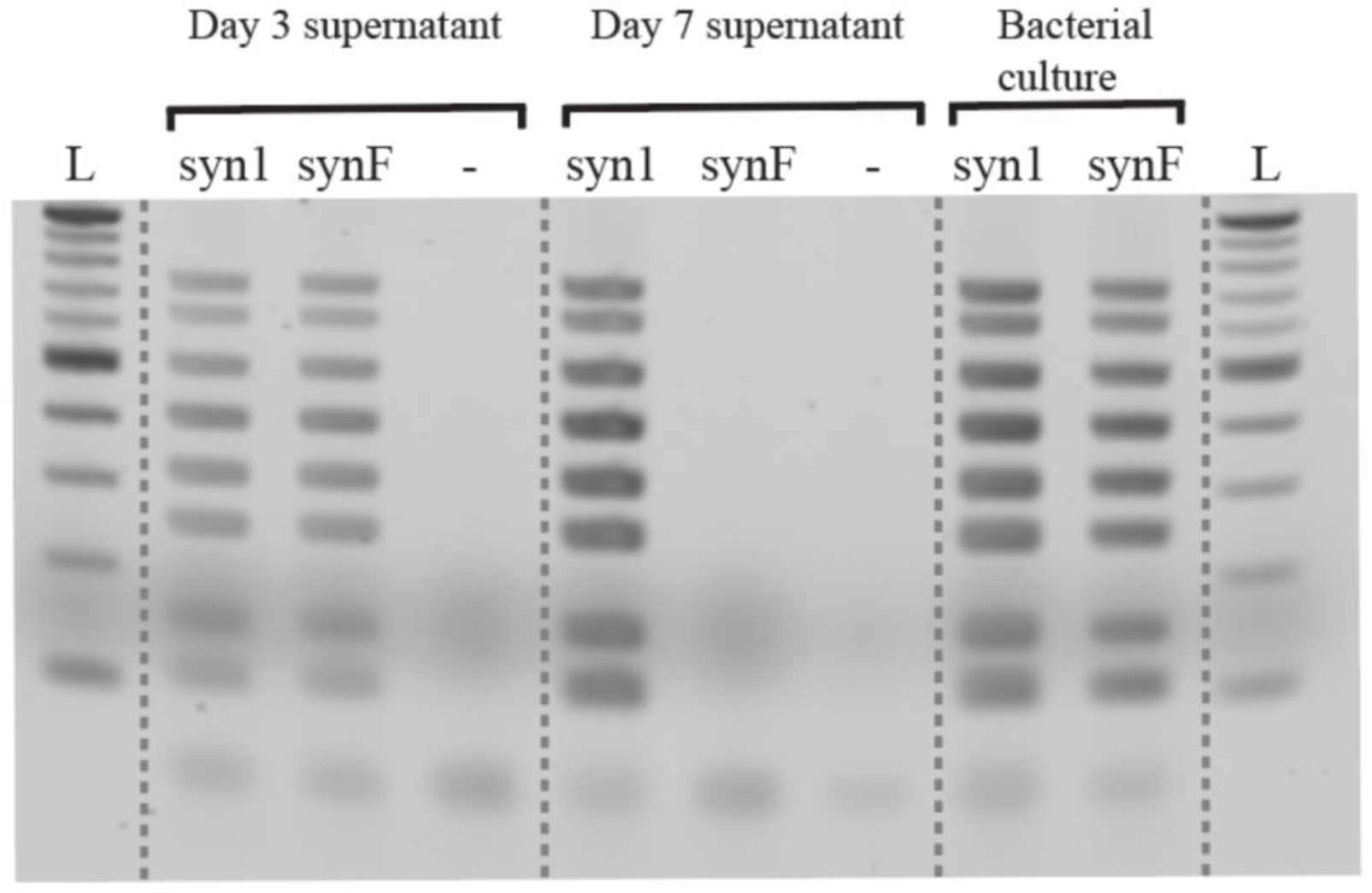
Detection of synthetic *Mmc* in HEK-293T cell cultures by multiplex PCR. Using eight sets of multiplex primers shown in Supplementary Table S3, PCRs were performed on the supernatant from cultures of HEK-293T cells inoculated with JCVI-syn1.0 (syn1), syn1.0F (synF), or a culture of HEK-293T cells without any bacteria (-). Actively growing bacterial cultures of each strain were used as positive controls. A molecular ladder (L) with markers every 100 bases from 100 bp to 1000 bp is shown in both outside lanes.

### Construction of JCVI-syn3B add-back mutants and CCU analysis after their incubation with mammalian cells

We identified a cluster of eight candidate genes (**Table 3**) that might be necessary for *Mmc* infection of mammalian cell lines. Although any or all of them might be necessary for synthetic *Mmc* survival in co-culture with mammalian cells, we hypothesized the putative lipoproteins MMSYN1_0179 and MMSYN1_0180, which are likely ABC transporter substrate binding proteins, and MMSYN1_0181, which is a membrane associated protein of unknown function, were logical candidates for enabling JCVI-syn1.0 binding to mammalian cells. Lipoproteins have been linked to interactions between mycoplasmas and eukaryotic cells, particularly with respect to adhesion (Pilo et al., 2007). In order to determine the contribution of each one of these genes to synthetic *Mmc* strain adherence and proliferation with cell hosts, we introduced them back to the genome of JCVI-syn3B at its dual *loxP* landing pad. We developed three different strains containing one of each of the target genes, MMSYN1_0179, MMSYN1_0180 or MMSYN1_0181. CCU analysis revealed that none of these genes alone enabled JCVI-syn3B survival for 10 days in co-culture with HEK-293T cells (**Figure S5**), confirming the suspicion that the presence of more than one of these genes might be necessary to recover the capacity of the minimal cell to infect host cells.

**Table 3.** Eight non-essential genes that might enable for *Mmc* survival in co-culture in mammalian cell lines. This cluster of eight genes was removed from JCVI-syn 1.0 to produce Syn1.0F. These genes are not present in JCVI-syn3A or JCVI-syn3B, except as in add back experiments using JCVI-syn3B.

| Start | End | Gene designation | Functional category | GenBank annotation |
| --- | --- | --- | --- | --- |
| 234726 | 236525 | MMSYN1_0179 | Cell envelope/<br>Transport &<br>binding<br>proteins | putative lipoprotein, possible<br>ABC sugar transporter<br>substrate binding protein |
| 236820 | 238610 | MMSYN1_0180 | Cell envelope/<br>Transport &<br>binding<br>proteins | putative lipoprotein, possible<br>ABC sugar transporter<br>substrate binding protein |
| 238692 | 239183 | MMSYN1_0181 | NULL | hypothetical protein (PFAM<br>analysis suggests it is part of<br>an ABC transporter) |
| 239198 | 241615 | MMSYN1_0182 | Transport &<br>binding<br>proteins | Pseudogene (malto-<br>saccharide ABC transporter,<br>permease; this<br>gene contains a frame shift<br>that may be the result of a<br>sequencing error; identified<br>by match to protein family<br>HMM PF00528) |
| 241630 | 244170 | MMSYN1_0183 | Transport &<br>binding<br>proteins | maltose ABC transporter<br>permease protein |
| 244172 | 245257 | MMSYN1_0184 | Transport &<br>binding<br>proteins | multiple sugar-binding<br>transport ATP-binding<br>protein MsmK |
| 245365 | 247662 | MMSYN1_0185 | Central<br>intermediary<br>metabolism | glycosyl hydrolase, family<br>65 |
| 247664 | 249268 | MMSYN1_0186 | Energy<br>metabolism | trehalose-6-phosphate<br>hydrolase (Alpha, alpha-<br>phosphotrehalase) |

To evaluate this hypothesis, we constructed two additional gene add-back strains, JCVI-syn3B::mCh-MMSYN1–0179-0181 and JCVI-syn3B::mCh-MMSYN1–0179-0186, comprising the cluster of the three genes we thought might enable mycoplasma survival evaluated in concert and the complete eight gene cluster, respectively. The gene MMSYN1_0182 is characterized as a pseudogene that produces a nonfunctional ABC transporter permease in JCVI-syn1.0 (**Table 3**). This gene was kept in the study to avoid possible transcription disruption of downstream genes and was not considered in further analysis. We also included a gene for the expression of mCherry fluorescent protein in the N-terminal region of both gene clusters, to verify cytoadherence to host cells with further fluorescence microscopy analysis.

In order to confirm the possible recovery of the survival phenotype by the synthetic *Mmc* mutants, we proceeded with the infectivity assay in HeLa cells, comparing the add-back mutant strains with JCVI-syn1.0::mCh and JCVI-syn3A::mCh. As mentioned above, JCVI-syn3A and JCVI-syn3B have the same genome, with the exception that JCVI-syn3B contains two *loxP* recombination sites for gene insertion. With the exception of JCVI-syn1.0::mCh, all mutant strains including JCVI-syn3A::mCh presented a negative color changing assay (this differs from a CCU assay in that only one dilution of each sample is tested and growth only indicates the presence of viable mycoplasmas) after 7 days post mixing with mammalian cells. Two out of three HeLa cell cultures inoculated with JCVI-syn3A::mCh and three out of three JCVI-syn3B::mCh-MMSYN1–0179-0181 inoculated into HeLa cells survived for 10 days. However, none of them was able to survive as long as JCVI-syn1.0::mCh, which could be detected up to 15 days post infection (**Figure 2A**). To validate this assay, we used PCR to detect *Mmc* 23S rRNA (species specific) and MMSYN1-0180 (strain specific) genes. As expected, the MMSYN1-0180 was detected only in the JCVI-syn1.0::mCh and JCVI-syn3B add-back mutants (**Figure 2B**).

**Figure 2.**
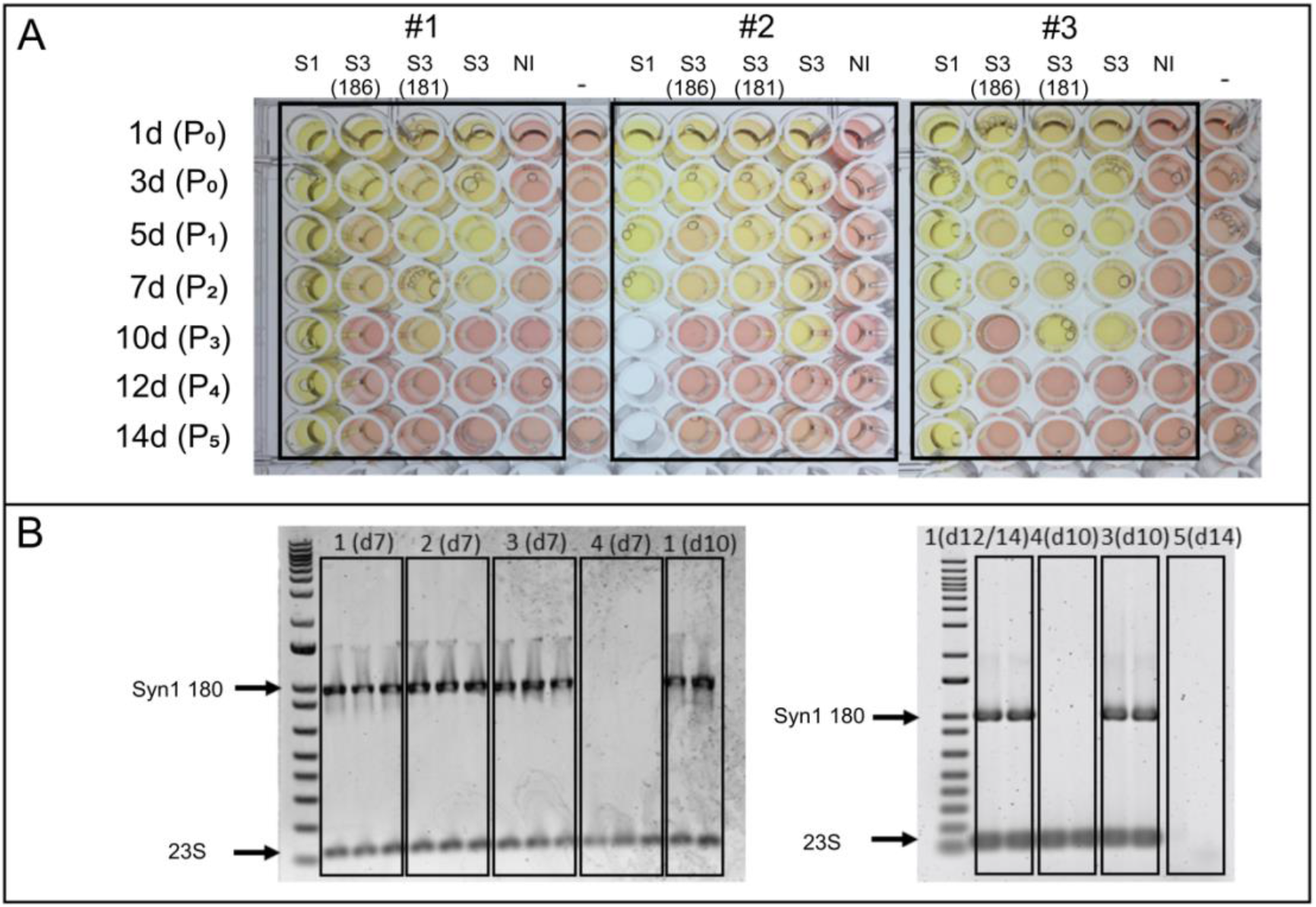
CCU assay of synthetic *Mmc* co-cultured with HEK-293T cells. (A) 100 μl of actively growing cultures of JCVI-syn1.0::mCh (S1), JCVI-Syn3B::MMSYN1–179-186 (S3(186)), JCVI-Syn3B::MMSYN1–179-181 (S3(181)), JCVI-syn3.0A::mCh (S3), or 100 μl of DMEM media (no infection) were added to cultures of HeLa cells (NI). At selected times over 14 days (d) a sample of supernatant was removed and a color changing assay was performed by adding 100 μl of the DMEM supernatant to 900 μl of SP4 mycoplasma growth media. These samples were incubated at 37□C for 7 days. The yellow color indicates there were viable cell in the sample because acid produced by bacterial metabolism altered the color of the phenol red pH indicator. Pure SP4 media is shown as a negative control to show the color without any bacterial growth (-). Passage number (P#) is given in parentheses. The analysis were performed in triplicate (#1, #2 and #3). (B) PCR was performed to detect *Mmc 23S rRNA* (23S, species specific) and *MMSYN1_0180* (Syn1 180, strain specific) genes in the supernatant from HeLa cell cultures inoculated with JCVI-syn1.0::mCh (1), JCVI-syn3B::MMSYN1–0179-0186 (2), JCVI-syn3B::MMSYN1–0179-0181 (3), JCVI-syn3.0A::mCh (4) or an uninoculated HeLa cell culture (5) at select time points (d) (obs., 1(d12/14), means sample JCVI-syn1.0::mCh at day 12 and 14). A molecular ladder (1 Kb Plus DNA Ladder, Invitrogen) is shown in outside lanes.

### Use of JCVI-syn3B::mCh-MMSYN1–0179-0181 and JCVI-syn3B::mCh-MMSYN1–0179-0186 add-back mutants for host-pathogen interaction studies

To verify the capacity of JCVI-syn3B::mCh-MMSYN1–0179-0181 and JCVI-syn3B::mCh-MMSYN1–0179-0186 mutants to adhere to mammalian cells, we analyzed the mycoplasmas co-cultured with HeLa cells using fluorescence microscopy, at 24 h and 48 h after infection (**Figure 3**). Although mycoplasma contamination cannot be determined using brightfield light microscopy, the mutant strains analyzed expressed the mCherry fluorescent protein, so we could employ epifluorescence microscopy to detect contaminating mycoplasmas in mammalian cell cultures. The HeLa cell cultures inoculated with minimized *Mmc* constructs were compared with cultures inoculated with JCVI-syn1.0::mCh and JCVI-syn3A::mCh. Uninfected HeLa cells were analyzed as negative control.

**Figure 3.**
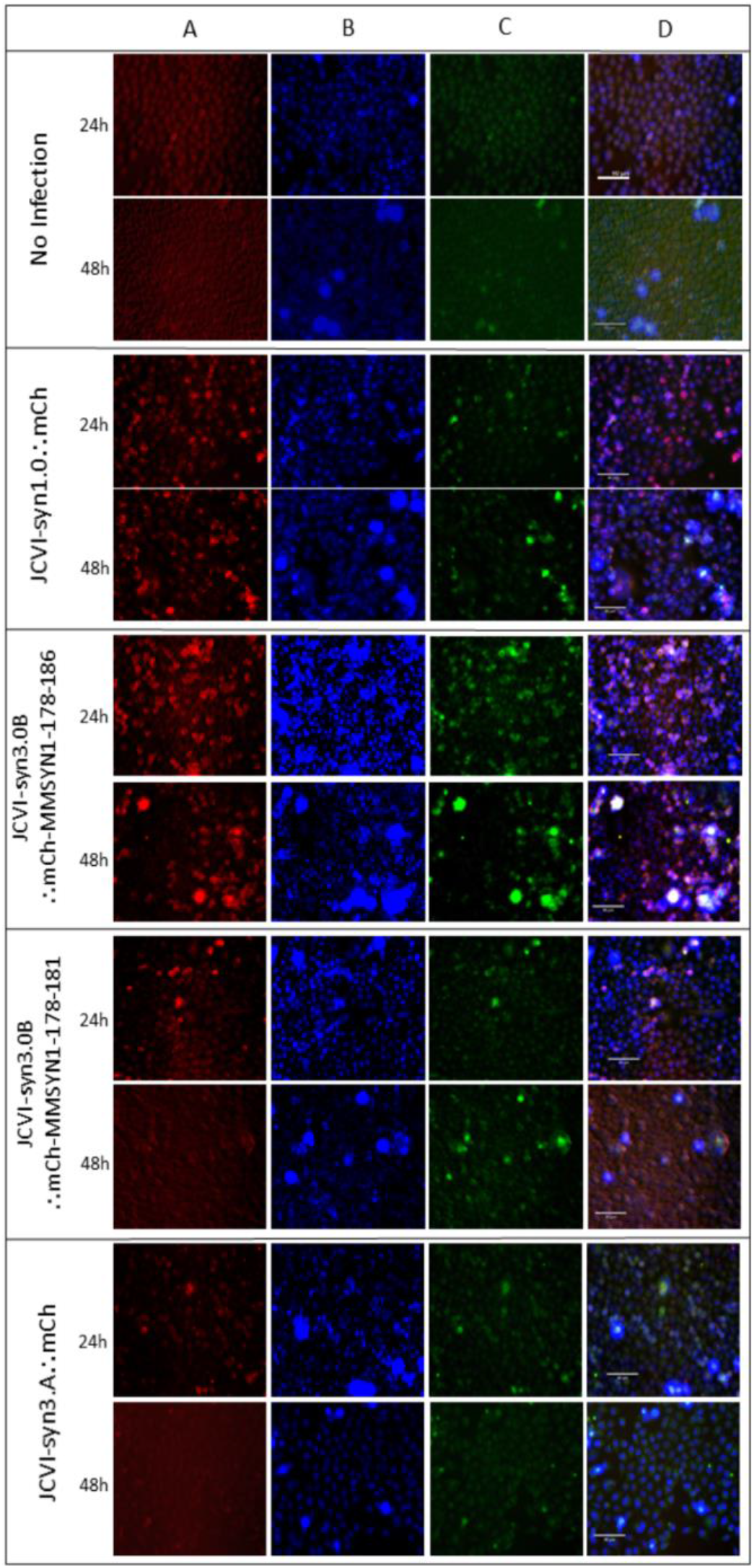
Live cell fluorescence microscopy. To verify the physical association of JCVI-Syn1.0::mCh, JCVI-Syn3A::mCh, JCVI-Syn3B::mCh-MMSYN1–179-181 and JCVI-Syn3B::mCh-MMSYN1–179-186 strains with viable mammalian cells, mycoplasmas co-cultured with HeLa cells were analyzed using fluorescence microscopy, at 24 h and 48 h post infection. This physical association of the mycoplasmas with the HeLa cells is likely adherence although it could be that some of the cells entered the HeLa cells. (A) mCherry fluorescence emission (red), (B) NucBlue fluoresce emission (blue), (C) CellROX fluoresce emission (green) and (D) merged fluorescence images. Scale bars indicate 90 μm.

In accordance with the CCU analysis, besides JCVI-syn1.0::mCh, none of the mutants were able to survive in HeLa cell culture more than 10 days. Even though, the live cell images showed the presence of mycoplasmas mutants cytoadhered to host cells at 24 h post infection. After 48 h, only JCVI-syn1.0::mCh and JCVI-syn3B::mCh-MSYN1–0179-0186 were still observed adhered to the mammalian cells (Figure 3A and Figure 4). While our assay could not distinguish whether the bacteria were adhering to the surfaces of the HeLa cells or internalized within the mammalian cells, our previous observation that in scanning electron micrographs JCVI-syn3B expressing the *U. parvum mba* gene were adhering to the surface suggests the cells here were adhering as well (Nishiumi *et al*., 2021).

**Figure 4.**
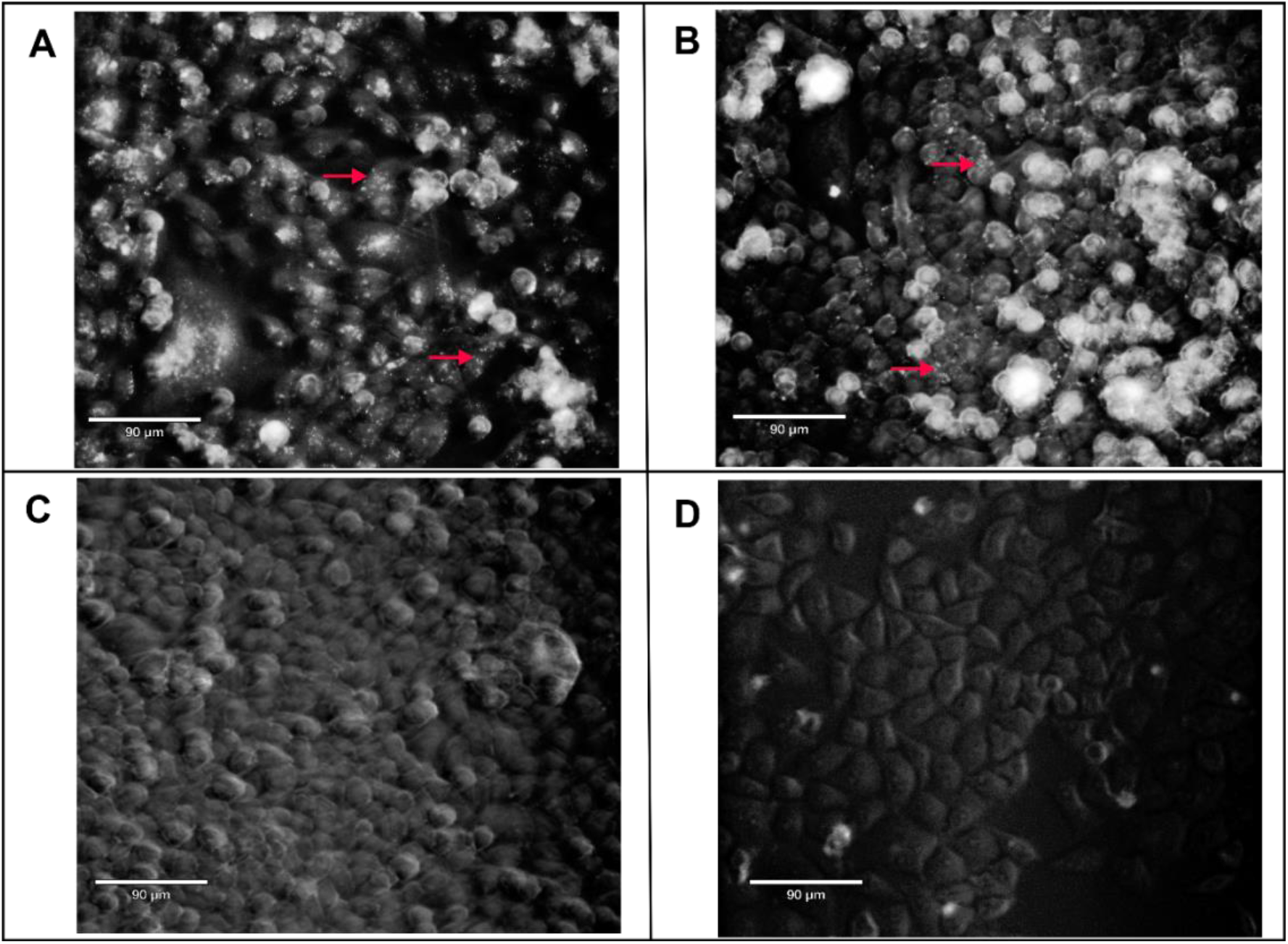
Live cell fluorescence microscopy of synthetic mycoplasmas co-cultured with HeLa cells for 48 h. Red arrows point to JCVI-Syn1.0::mCh (small white dots) (A) and JCVI-syn3B::mCh-MMSYN1–179-186 (B). Samples (C) and (D) showed no presence of JCVI-syn3B::mCh-MMSYN1–179-181 or JCVI-syn3A::mCh after 48 h of co-culture with HeLa cells, respectively.

The production of overall reactive oxygen species (ROS) in HeLa cells was also monitored by a probe, CellROX, 24 h and 48 h after synthetic *Mmc* infection (**Figure 3C**). Production of ROS was clearly visible in JCVI-syn1.0::mCh and JCVI-Syn3::mCh-MMSYN1–0179-0186 infected HeLa cells, in comparison with the other analyzed strains. No ROS was observed in the uninfected HeLa cells. **Figure 3B** shows NucBlue bound to HeLa cell DNA, and **Figure 3D** shows merged fluorescence images. The results indicated that the presence of the candidate eight gene cluster in the synthetic *Mmc* strains induced oxidative stress in HeLa cells, compared to cells inoculated with JCVI-syn3A::mCh and uninfected cells.

### Phagocytic activity of dHL-60 cells in the presence of synthetic Mmc strains

After showing minimized *Mmc* was incapable of parasitizing mammalian cells, we hypothesized these cells, which lacked most of the proteins on the surface of wild type *Mmc*, might be invisible to arms of the human immune system. We also hoped to probe mechanisms of *Mmc* interactions with their hosts in natural infections. To test this, we mixed fluorescently labelled synthetic *Mmc* strains with neutrophil-like, undifferentiated and differentiated human promyelocytic leukemia cells, HL-60 and dHL-60, respectively. We analyzed the reactions in these mixtures using flow cytometry. We also wanted to know if presence of our candidate genes for enabling *Mmc* adherence to mammalian cells would alter the action of the phagocytizing cells on the minimized mycoplasma cells.

First, we analyzed if the strains JCVI-syn1.0::mCh and JCVI-syn3A::mCh were capable of inducing a phagocytic activity in dHL-60 cells (**Videos S1 and S2**, respectively). Using a 3D Cell Explorer Microscope (Nanolive), we performed live cell imaging at 37°C. The formation of dHL-60 cell membrane projections (pseudopodia) and neutrophil extracellular traps (NET) (Scieszka et al., 2020; Thiam et al., 2020), by the dHL-60 cells towards JCVI-syn1.0::mCh were visible. In support of our hypothesis, JCVI-syn3A::mCh did not induce a strong immunogenic response in dHL-60 cells in comparison with JCVI-syn1.0::mCh.

Flow cytometry analyses agreed with previous observations and indicated a higher phagocytic activity by dHL-60 cells in the presence of JCVI-syn1.0::mCh, rather than JCVI-syn3A::mCh. HL-60 cells were used as negative controls for the phagocytic activity. By simultaneous measurement of cellular light scatter and fluorescence, extracellular bacteria, phagocytes, and non-phagocytes could be discriminated and PI quantified (**Figure S6**). Phagocytosis was evaluated after 30, 150 and 210 min following incubation of synthetic *Mmc* cells with neutrophils. The mean PI values found for HL-60 cells were statistically lower than dHL-60 at all time points, with exception of PI values induced by JCVI-syn3A::mCh, which HL-60 and dHL-60 cell’s PI did not present any statistical difference (**Figure 5**).

**Figure 5.**
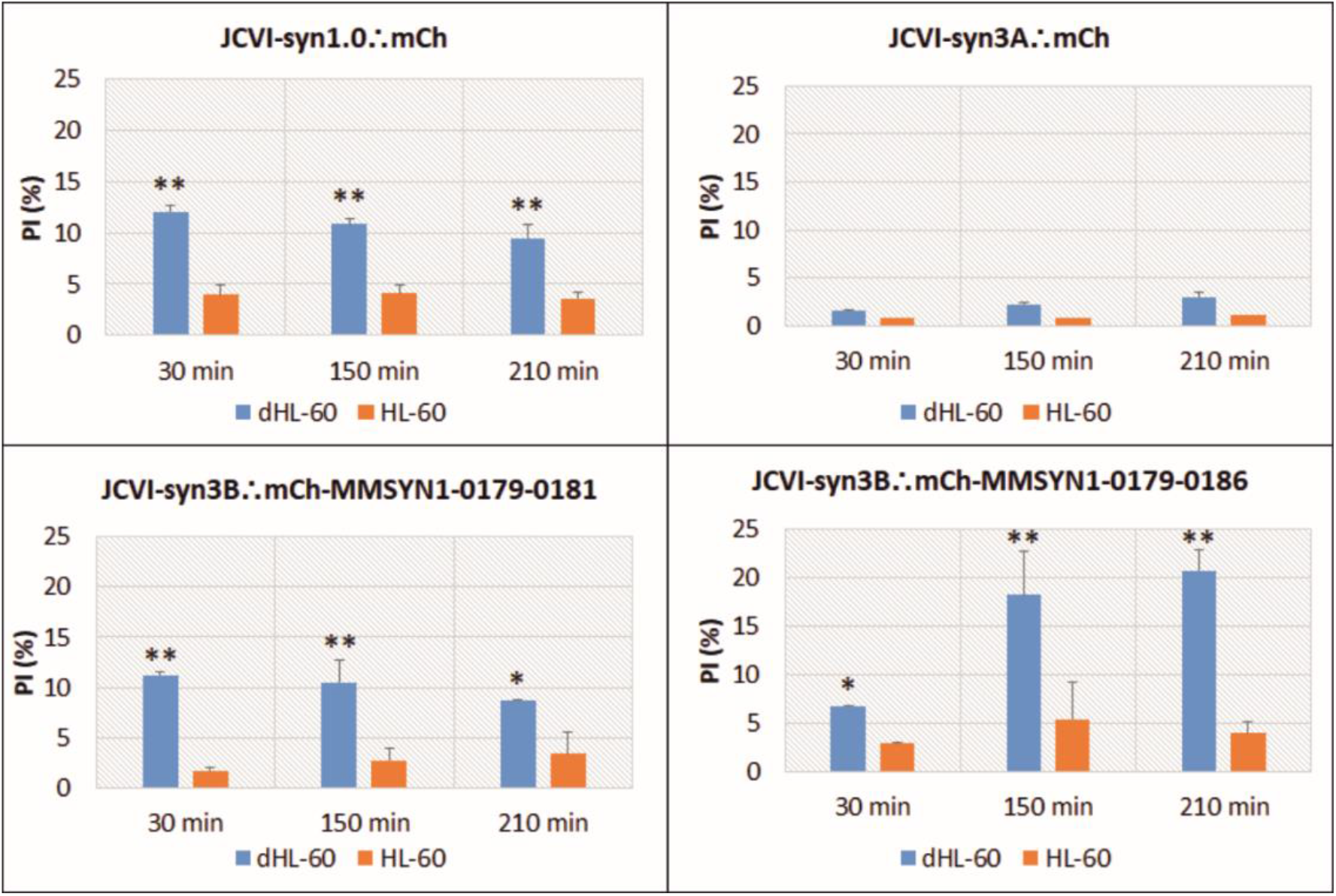
Phagocytic Index (PI) of HL-60 and dHL-60 neutrophil cells, in the presence of synthetic mycoplasma strains. With the exception of JCVI-syn3A::mCh, the difference between the PI means of dHL-60 versus HL-60 (control) cells were statistically significant. Data are presented as means ± SD standard error of two observations for each synthetic cell and time point. * P < 0.01, ** P < 0.001 versus control.

According to the statistical analysis, the Y ~ B * T + (1|S) model (4^th^ model) was the best representation of our data set (**Table S4**), showing that synthetic *Mmc* cell type and time of inoculation were both responsible for the phagocytic activity of dHL-60, with a total variance of 48% within population. However, only JCVI-syn3B::mCh-MMSYN1-0179-0186 was able to induce a statistically higher phagocytic activity in dHL-60 over time. Pointing to a major role of synthetic *Mmc* cell type in inducing phagocytosis rather than time. No PI variance was observed for all HL-60 treatments over time.

Comparison between strains showed that the phagocytic ability of dHL-60 cells (**Figure 6**) was the lowest in the presence of JCVI-syn3A::mCh. Interestingly, no significant difference was observed between the PIs of dHL-60’s co-culture with JCVI-syn1::mCh and JCVI-syn3B::mCh-MMSYN1-0179-0181, in all incubation times. However, the PI values induced by JCVI-syn3B::mCh-MMSYN1-0179-0186 were statistically higher at 150 and 210 min after inoculation with dHL-60 cells (F**igure S7**).

**Figure 6.**
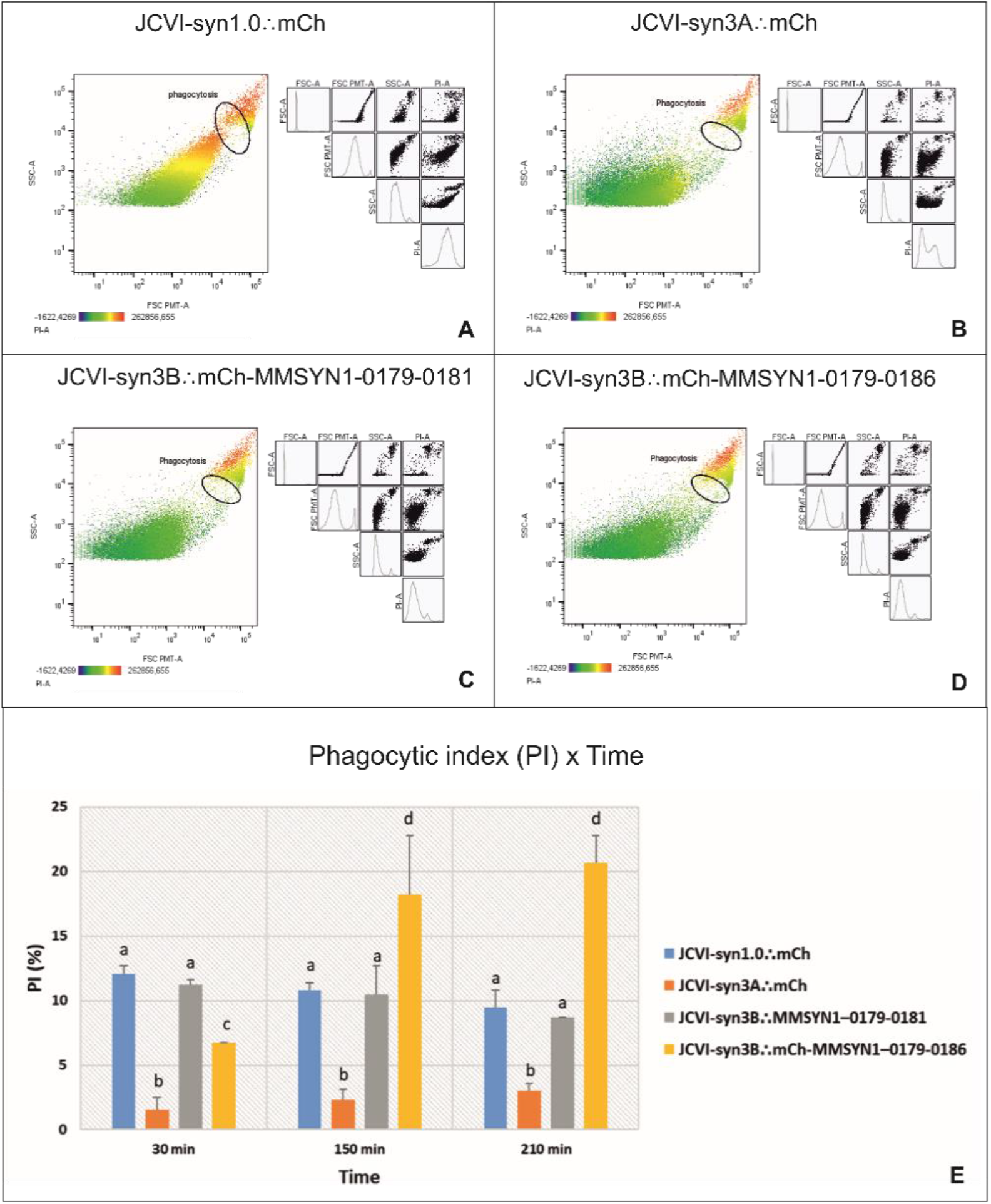
Flow cytometry analysis of dHL-60 neutrophils cells, in the presence of synthetic *Mmc* strains, analyzed at 30 min, 150 min and 210 min after mixing the two cell types. (A-D) Graphs of representative flow cytometry plots (FSC PMT-A X SSC-A). (E) Phagocytic Index (PI) of dHL-60 presented as mean ± standard error of two independent experiments over time (30 min, 90 min, 210 min). For all variables with the same letter, the difference between the means is not statistically significant (p <0.01).

## Discussion

Because mollicutes are parasitic bacteria that lack cell walls, their cell membrane is the surface of interaction between bacterium and host cells. These interactions enable mycoplasmas to obtain essential nutrients and also are critical to mycoplasma pathogenicity. Multiple proteins, such as adhesins, lipoproteins and even common housekeeping enzymes, have been identified on the cell surface of various mycoplasmas species that are important if not essential for colonization of their hosts (Razin et al., 1998). Nevertheless, our understanding of the specificity and pathogenic mechanisms of the bacteria comprising the *Mycoides* group of mycoplasmas, which is known to cause economically significant diseases in ruminants, is far from complete.

Using an assay that detected living mycoplasma cells that survive in co-culture with mammalian cells, we confirmed our previous result showing that our minimized *Mmc* cells did not survive more than 7-10 days in co-culture with Hela or HEK-293T cells (Nishiumi *et al*., 2021). There was no difference in the capacities of HEK-293T cells and HeLa cells to support proliferation of synthetic *Mmc* strain JCVI-syn1.0 (**Table 2**), indicating that the genes necessary for the survival phenotype were not specific for either mammalian cell line. Further, the secreted products from either cell line were not sufficient for mycoplasma survival in conditioned medium.

That the minimized *Mmc* is unable to survive in co-culture with mammalian cells more than a few days is not surprising. The minimization process resulted in a cell containing less than half the number of genes present in wild type *Mmc*, and perhaps more importantly in this circumstance, deletion of 71 of 87 lipoprotein encoding genes. Thus, a huge fraction of the proteins that decorate the surface of *Mmc* and potentially mediate host-parasite interactions are not in the minimized genomes.

Comparing the infectivity assay results from JCVI-syn1.0, JCVI-syn3.0 and several intermediate minimized genome strains, we identified the cluster of genes that might be correlated with the capacity of certain synthetic *Mmc* strains to survive when co-cultured with mammalian cells. We hypothesized that adding back the candidate genes, or at least three of them, to the minimal cell JCVI-syn3B, which lacks the capacity to adhere to and grow in mammalian cell co-culture, would also restore its ability to infect cultured mammalian cells. However, none of the mutants were able to survive with the host; although, JCVI-syn3A::mCh and JCVI-syn3B::MMSYN1–0179-0181 were capable of surviving a few days longer than the minimized strains JCVI-syn2.0 and JCVI-syn3.0.

Our microscopy data indicated survival equates with attachment to (or possibly internalization by) the mammalian cells (internalization of *Mmc* in non-phagocytic cells has been demonstrated (Di Teodoro *et al*., 2018)). Similar to JCVI-syn1.0::mCh, the addition of the candidate genes improved the cytoadherence of the minimal cell and after 48 h of inoculation only JCVI-syn3B::mCh-MMSYN1–0179-0186 was still observed adhered (or less probably internalized by) with HeLa cells (**Figure 3**). The presence of MMSYN1–0179-0186 genes may have conferred a tighter interaction between the synthetic mycoplasmas and the host cells, compared to JCVI-syn3A::mCh and JCVI-syn3B::MMSYN1–0179-0181 cells. As shown in Table 2, the Reduced Genome Design strain RGD1.0-2 was the only strain incapable of surviving 7 days in co-culture with HeLa and HEK-293T cells. Our data showed that the genes MMSYN1–0179-0186 enabled attachment but not proliferation. If the genes that necessary for mycoplasma growth were present in any of the 1/8^th^ genome segments other than number 2, then any of the 7 other RGD1.0 strains lacking one of those genes would likely have not grown in the assay shown in Table 2. That was not the case, so the genes necessary for parasitism of the mammalian cells are likely in genome segment 2. An examination of the 40 genes deleted from that segment (other than MMSYN1–0179-0186) (Hutchison *et al*., 2016) does not suggest any obvious candidates.

The experiments presented here that showed attachment rather than proliferation are consistent with our findings in an earlier study where we installed the *Ureaplasma parvum* serovar 3 major surface antigen *mba* in the JCVI-syn3B landing pad. In that study, the *mba* expressing JCVI-syn3B mutant was able to bind to Hela cells, but not proliferate. It was the addition of a second pathogenicity related *U. parvum* gene, *UpVF* that appears to enable the cells to proliferate in co-culture with HeLa cells (Nishiumi *et al*., 2021). Presumably, there are non-essential *Mmc* genes that enable cell proliferation once the cell has attached to the mammalian cells using proteins expressed from the MMSYN1–0179 to MMSYN1-0186 gene set. On the other hand, the infectivity assay of JCVI-syn1.0F, a strain derived from JCVI-syn1.0 lacking only MMSYN1-0179-0186 gene cluster, showed that this strain completely lost its capacity to survive in co-culture with and infect mammalian cell cultures. Although those genes were not capable of making JCVI-syn3B infectious again they can be considered essential for the maintenance of synthetic *Mmc* capacity to interact with the cell host and might be likely required for survival of *Mmc in vivo*.

The genes MMSYN1-0179 and MMSYN1-0180 are lipoproteins, which are well known to have a central role in interactions between mycoplasmas and eukaryotic cells (Browning et al., 2011; Yiwen et al., 2021). The genes MMSYN1-0183 and MMSYN1-0184 are annotated as ABC transporter proteins, and MMSYN1-0185 and MMSYN1-0186 are both related with cell metabolism. The substrate(s) of these transporters is/are unknown. The MMSYN1-0181 gene is the only one of these proteins whose function is unknown (**Table 3**). Many Gram-positive bacterial lipoproteins are substrate-binding proteins of ABC transporter systems responsible for the acquisition of multiple nutrients including amino acids and short peptides, sugars, polyamines, and many metal ions (Nguyen et al., 2020). It is also known that acquisition and metabolism of carbohydrates are essential for host colonization and pathogenesis of bacterial pathogens (Davies et al., 2021; Tan et al., 2015). Since the MMSYN1-0180-0186 genes are present in the same operon (MMSYN1-0179 is transcribed by itself, data not shown), they might function together contributing with the synthetic cell adherence to the cell host. Interestingly, a predicted maltose ABC transporter *malF* was found to be necessary for the persistence of *Mycoplasma gallisepticum* in infected birds (Tseng et al., 2013). This datum also agrees with our observation that the putative maltose ABC transporter permease MMSYN1_0183 might play an important role in *Mmc* cytoadherence .

During proliferation in the host, successfully surviving mycoplasmas generate numerous metabolites, including hydrogen peroxide, ammonia and hydrogen sulfide (Yiwen *et al*., 2021). Absence of genes involved in hydrogen peroxide synthesis from the minimized *Mmc* strains precludes them from expression of ROS (Breuer *et al*., 2019). In our experiments JCVI-syn1.0::mCh and JCVI-syn3B add-back mutants induced oxidative stress in HeLa cells. The source of the oxidative stress may have been increased ROS generation, or cellular antioxidant capacity induction in mammalian cells due to the attached mycoplasma cells (or a combination of both) (Ji et al., 2019). When attached to the surface of eukaryotic cells mycoplasmas can interfere and alter cellular pathways and the host organism engages upon infection a series of responses that involves a number of signaling pathways, eventually resulting in the activation of the immune system (Benedetti *et al*., 2020).

In order to quantify the capacity of synthetic *Mmc* strains to solely induce an immunological response in dHL-60 cells, we performed a phagocytic assay. Neutrophils provide the first line of defense of the innate immune system in the control of common bacterial infections. They recognize particulate substrates of microbial origin and sequester the cargo via phagocytosis and/or NET formation (NETosis) (Manfredi et al., 2018; Rosales, 2020; Scieszka *et al*., 2020; Thiam *et al*., 2020). NETs are web-like DNA structures decorated with histones and cytotoxic proteins that are released by activated neutrophils to trap and neutralize pathogens during the innate immune response (Thiam *et al*., 2020). Agreeing with experiments using *Mycoplasma agalactiae* (Cacciotto et al., 2016), we observed NET formation by dHL-60 in co-culture with JCVI-syn1.0::mCh using a 3D Cell Explorer Microscope (**Video S1**).

Mycoplasma lipoproteins are also considered the primary proinflammatory substances and are capable to interact not only with epithelial cells but also to the leukocytes of the host organism (Borchsenius et al., 2020; Medzhitov, 2007; Shimizu, 2016; Yiwen *et al*., 2021). Accordingly, the variance observed in PI of dHL-60 cells was significantly related to the bacteria genotype. We accounted for non-specific binding of the microbes to the host cell membranes by including HL-60 cells as controls to our dHL-60 experiments (**Figure 5**). With the exception of JCVI-syn3A::mCh, all synthetic *Mmc* strains were capable to induce phagocytosis in dHL-60 cells, indicating that the presence of the genes MMSYN1-0179 up to MMSYN1-0181 or up to MMSYN1-0186 into JCVI-syn3B restored the capacity of the minimal cell to promote phagocytic activity in neutrophils.

Surprisingly, dHL-60 showed the highest level of PI when co-cultured with JCVI-syn3B::mCh-MMSYN1-0179-0186, in comparison with JCVI-syn1.0::mCh and JCVI-syn3B::mCh-MMSYN1-0179-0181, which did not present statistical difference between strains and over time (**Figure 6**). Although the host immune system can eradicate the invading mycoplasma in most cases, it is known that mycoplasmas can employ a series of immune escape strategies to ensure their continued parasitism of their hosts. For instance, capsular polysaccharides and invasive enzymes are crucial for anti-phagocytosis and immunomodulation (41). Furthermore, bacterial cell walls, which are absent in mycoplasmas, are important triggers of mammalian antibacterial innate immune responses. This might further reduce the phagocytosis inducing ability of mycoplasmas. Differently from JCVI-syn3A::mCh, such characteristics might be also present in JCVI-syn1.0, which genome resembles the one from the natural *Mmc*, explaining the higher capacity of JCVI-syn3B::mCh-MMSYN1-0179-0186 to induce phagocytosis in dHL-60. This result indicates that MMSYN1-0179-0186 genes might be excellent candidates for the development of new vaccines.

Mycoplasma infection is likely the result of a number of actions involving several surface membrane components, allowing the pathogen to adhere tightly to the host cell surface, obtain needed metabolites from the host cell, and proliferate. The JCVI-syn3.0 minimized genome is roughly half the size of the synthetic cell JCVI-syn1.0 genome (531 kbp and 1079 Kbp, respectively). For instance, the *Mmc* GM12 genome encodes for 87 lipoproteins, while the minimal cell produces only 15 (Hutchison *et al*.,2016). Although adding back the *MMSYN1-0179-0186* gene cluster into JCVI-syn3B induced an immunological response in neutrophils, it was not enough to restore the full capacity of the minimal cell to infect a mammalian cell culture.

Indeed, a recent study produced the mutant *Mmc* strain GM12::YCpMmyc1.1-Δ68 with complete abolishment of pathogenicity (Jores et al., 2019). Animals infected with the *Mmc* GM12 strain developed specific clinical signs (fever, heavy breathing, septicemia, etc.) and were all euthanized by 6 days post infection. On the contrary, the goats infected with GM12::YCpMmyc1.1-Δ68 did not develop such signs and were healthy for the entire course of the experimentation. For the production of this strain 10% of the genome content were removed, comprising 68 genes from different functional categories, although the MMSYN1-0179-0186 cluster was not removed. This observation points towards a regulatory complexity of *Mmc* infection.

However, the inability of JCVI-syn3A::mCh to induce an immunological response in neutrophils, and its possible invisibility to the human immune system, together with its inability on its own to infect mammalian cells or its natural host, goats, indicates that the process of genome minimization might contribute with the development of synthetic cells to function as specific drug delivery systems. Consider that the therapeutic capacity of most cell-based anti-cancer using patient derived cells is limited to the <10kb of DNA that can be delivered to these cells using lentivirus or retrovirus vectors. A minimized *Mmc* cell would be capable of carrying cancer fighting payloads of up to or more than 500 kb. The cost of producing therapeutic minimized mycoplasmas tailored for specific cancers would likely be vastly lower than the current cell-based therapies.

This report not only advance the knowledge of underlying molecular and infection mechanisms in a pathogenic specie that constitute the *M. mycoides* cluster of mycoplasmas, but also confirms that the minimized synthetic *Mmc* strains JCVI-syn2.0, JCVI-syn3.0 and its derivatives lack the cellular machinery necessary for infecting and contaminating mammalian cell cultures. If these strains are to be used as a potential model to blueprint the genes necessary for life, the extra precautions for limiting their use such as requiring work be done in Biosafety Level 2 laboratories are no longer necessary. Nevertheless, further evaluation is necessary to determine which specific gene(s) is (are) still necessary to confer the survival phenotype in minimized *Mmc* synthetic strains.

The possibility to use safe laboratory synthetic mycoplasma strains for research purposes opens the opportunity to identify the actual role of suggested virulence determinants in mycoplasma and, consequently, to better understand the mechanisms that drives host–pathogen interactions. Furthermore, the reduced genome synthetic cell might also serve as a platform to test known and potential virulence factors from different bacteria, whose basis for interaction might be confounded by multiple factors expressed from their large genome, and contribute with the design of new potential antimicrobials and vaccines.

### Limitations of the study

We confirmed previous findings that the minimized synthetic cells that contain subsets of the *Mmc* genome were unable to parasitize cultured mammalian cells. New work identified a cluster of eight non-essential genes (MMSYN1-0179-0186) mediated *Mmc* attachment to mammalian cells, but did not enable parasitism of mammalian cells. In future experiments we hope to use electron microscopy to visualize the interface between the minimized *Mmc* cells expressing MMSYN1-0179-0186 and mammalian cells. We will compare that with the interface of wild type mammalian cells to try to gain insight into how the MMSYN1-0179-0186 proteins mediate attachment and how mycoplasma attachment and effective parasitism may differ. The inability of the minimized *Mmc* to infect mammalian cell cultures and the poor phagocytization of the minimized *Mmc* by human neutrophil-like cells suggests these cells are to some extent invisible to the mammalian innate immune system. In future work we will investigate this more thoroughly by infecting animals with the minimized *Mmc* and determining whether other phagocytic cells attack the minimized *Mmc*.

## Data and code availability

All data reported in this paper will be shared by the lead contact upon request.

This paper does not report original code.

Any additional information required to reanalyze the data reported in this paper is available from the lead contact upon request.

## Acknowledgments

This study was supported by Brazilian Agricultural Research Corporation (EMBRAPA, Brazil, #21195.002926/2019-98) and National Institute of Science and Technology in Synthetic Biology (INCT BioSyn - CNPq, Brazil, #465603/2014-9), and by the United States National Science Foundation grants MCB 1840320, MCB 1818344 and MCB 1840301. The authors thank Kim Wise for his valuable comments and David Bianchi for his assistance evaluating the functions of *Mmc* non-essential genes MMSYN1_0179-0186.

## Author contributions

DMCB and DMB performed research, analyzed data, and wrote the paper; NAG, LS and MRL performed research and analyzed data; LAMPM analyzed data; MF and JIG designed research, analyzed data, and wrote the paper.

## Declaration of interests

The authors declare no competing interests.

## STAR★Methods

### Key resources table

| REAGENT or RESOURCE | SOURCE | IDENTIFIER |
| --- | --- | --- |
| Bacterial and virus strains |  |  |
| NEB 5-alpha Competent *Escherichia coli* | New England BioLabs | C2987H |
| Chemicals, peptides, and recombinant proteins |  |  |
| PrimeSTAR Max DNA Polymerase | Takara | R045B |
| OneTaq® Quick-Load® DNA Polymerase | New England BioLabs | M0509S |
| DpnI | New England BioLabs | R0176S |
| Critical commercial assays |  |  |
| Gibson Assembly® Master Mix | New England BioLabs | E2611L |
| Zyppy Plasmid Miniprep Kit | Zymo Research | D4019 |
| DNA Clean & Concentrator-5 | Zymo Research | D4013 |
| CellROX® Green Reagent | ThermoFischer | C10444 |
| NucBlue™ Live ReadyProbes™ Reagent (Hoechst 33342) | ThermoFischer | R37605 |
| Live Cell Imaging Solution | Invitrogen | A14291DJ |
| Experimental models: Cell lines |  |  |
| HeLa | ATCC | ATCC CCL-2 |
| HEK-293 | ATCC | ATCC CRL-1573 |
| HL-60 | ATCC | ATCC CCL-240 |
| Experimental models: Organisms/strains |  |  |
| JCVI-syn1.0 | Gibson et al., 2010 | GenBank: CP002027 |
| JCVI-syn2.0 | Hutchison et al., 2016 | GenBank: CP014992.1 |
| JCVI-syn3.0 | Hutchison et al., 2016 | GenBank: CP014940.1 |
| JCVI-syn3A | Breuer et al., 2019 | GenBank: CP016816.2 |
| JCVI-syn3B | Pelletier et al., 2021 | N/A |
| JCVI-syn1.0F | Gibson et al., 2010 | N/A |
| RGD1.0-1 through RGD1.0-8 | Hutchison et al., 2016 | N/A |
| JCVI-syn1.0::mCh | Pelletier et al., 2021 | N/A |
| JCVI-syn3.0::mCh | Pelletier et al., 2021 | N/A |
| JCVI-syn3B::mCh-MMSYN1–0179-0181 | This paper | N/A |
| JCVI-syn3B::mCh-MMSYN1–0179-0186 | This paper | N/A |
| Oligonucleotides |  |  |
| For all primers used in this study | See Tables S2 and S3 | N/A |
| Recombinant DNA |  |  |
| PRS313 | ATCC | 77142 |
| Pmod2-loxpurolox-sp-cre | Pelletier et al., 2021 | GenBank: MN982903.1 |
| Plasmids constructed in this study | See Figure S1 | N/A |
| Software and algorithms |  |  |
| FlowJo v10.6.1 | BD Biosciences | https://www.flowjo.com/solutions/flowjo/downloads |
| R | R Core Team, 2019 | https://www.r-project.org/ |
| Other |  |  |
| BD FACSAria™ II flow cytometer | BD Biosciences | 657590 |
| 3D Cell Explorer-fluo | Nanolive | Model CX-F |
| Revolve Fluorescence Microscope | Echo Laboratories Inc | R4 |

### Resource availability

#### Lead contact

Further information and requests for resources should be directed to and will be fulfilled by the Lead Contact, John Glass.

#### Materials availability

Materials generated in this study are available from the Lead Contact with a completed Material Transfer Agreement.

### Experimental model and subject details

#### Mycoplasma strains and mammalian cells cultures

With the exception of JCVI-syn3B (GenBank accession number acquisition in progress), all the synthetic mycoplasma strains used were previously described (Gibson *et al*., 2010; Hutchison *et al*., 2016; Pelletier *et al*., 2021). Similar to JCVI-syn3A, JVCI-syn3B possesses a second rRNA operon copy, lacks a gene (*MMSYN1_0531*) coding for an efflux protein, and has 16 protein coding genes that were added back into the JCVI-syn3.0 genome to render the cell with a more wild type *Mmc* morphology and growth rate (Breuer *et al*., 2019; Pelletier *et al*., 2021). The strain also contains a landing pad system (dual LoxP sites) that facilitates genetic manipulation. The genome sizes of each strain used in this work can be found at **Table S1**.

We maintained synthetic *Mmc* strains in SP4 medium containing a phenol red preparation supplemented with 17% (v/v) KnockOut™ Serum Replacement (Gibco) and 3 μg/ml tetracycline (Tet) (Williamson and Whitcomb, 1975). Cultures of the mCherry (mCh) expressing strains were grown in SP4 with no Tet. All strains were cultivated in static growth chamber at 37 °C.

HEK-293T and HeLa cells (ATCC®) were maintained in Dulbecco’s modified Eagle’s medium (DMEM) (Gibco) with 10% fetal bovine serum, 100 units/mL of penicillin, 100 μg/mL of streptomycin, and 0.25 μg/mL Amphotericin B at 37°C and 5% CO_2_.

The HL-60 promyelocytic cell line (ATCC®CCL-240™promyeloblast human cell line) was cultured in Iscove’s Modified Dulbecco’s Medium (IMDM) (Gibco) supplemented with 10% (v/v) FBS (Sigma) and 1% penicillin (Gibco) at 37°C with 5% CO_2_. To exhibit phagocytic activity and responsiveness to chemotactic stimuli the undifferentiated HL-60 cells were differentiated (dHL-60) by culture in Roswell Park Memorial Institute (RPMI) 1640 Medium (Gibco), supplemented with 10% (v/v) FBS (Sigma) and 1% penicillin (Gibco), in the presence of 1.25% (v/v) dimethyl sulfoxide (ATCC) for 5 days prior to use (Collins et al., 1978).

### Method details

#### Infectivity assay

The synthetic *Mmc* strains were grown in SP4 medium and once cultures had entered a stationary growth phase as indicated by acidification of the media causing the phenol red to turn orange (about 20 h), 100 μl of each sample was added to 5 ml of media in a mammalian monolayer cell culture (approximately 30-50% confluent) growing in a T25 25 cm^2^ flask. 100 μl of DMEM media were used to inoculate a culture of HeLa cells as negative control (no infection). The infected mammalian cells were incubated at 37°C and 5% CO_2_. Cultures were grown for 15 days and passaged (culture medium changed) each 2-3 days of cultivation. Once the mammalian cell culture reached confluence (~7 days) the culture was the cell monolayer was released from its attachment surface by trypsinization, and 10% of the cells were mixed with 9 volumes of DMEM in a new flask. The medium was removed, and 2 ml of Dulbecco’s phosphate-buffered saline (DPBS, Gibco) were used to wash the cells. DPBS was removed and 1 ml of trypsin 2.5% (ThermoFisher Scientific) was added to the culture flask. Mammalian cells were allowed to incubate at 37°C until they detached from the surface. Once trypsinization was complete 1 ml of culture medium was added and the cells half were transferred into a fresh 25T flask by adding 500 μl of cell suspension into 4.5 ml fresh medium.

To ensure that mycoplasmas could not grow in the mammalian cell medium (DMEM + 10% FBS), fresh and conditioned media only were also inoculated with each strain then incubated at 37°C for 7 days. HeLa and HEK-293T cells were grown for 2 days in fresh DMEM + 10% FBS, media was removed from the culture and filtered sterilized to produce the conditioned media.

To verify the presence of mycoplasma in the mammalian cell culture the determination of color changing unit (CCU) assay was performed at selected time points during the 15 days of co-cultivation.

#### Determination of Color Changing Unit (CCU)

To titrate mycoplasmas, specimens were diluted in serial 10-fold steps in SP4 mycoplasma liquid medium, which contained phenol red as a pH indicator. As mentioned before, bacterial metabolism leads to a pH shift that in turn causes the phenol red indicator to change color from red to orange and then to yellow indicating cell growth. The highest dilution that produced a color change on incubation at 37°C for 7 days was the end point of the titration and was considered to contain 1 color changing unit (Cherry and Taylor-Robinson, 1970).

Our CCU assays were performed in SP4 medium supplemented with 3 μg/ml Tet, with the exception of mCherry expressing strains that were cultivated in SP4 without Tet. Briefly, 100 μl of the growing culture was added to 900 μl fresh SP4 media followed or not by serial 10-fold dilutions as described previously (Benders et al., 2010; Dedieu and Balcer-Rodrigues, 2006; Lartigue *et al*., 2009). CCU assays were incubated at 37°C for 1 week.

Single or multiplex PCRs were also performed to confirm the presence of the synthetic *Mmc* strain at CCU samples. 2 μl of the culture medium was used as template. The PCR conditions were 94°C 30 sec; 30 cycles of 94°C for 30 s, 52°C for 30 s, and 68°C for 2.5 min; and, followed by 68°C for 5 min. The conditions for multiplex PCR were 94 °C for 15 min, then 35 cycles of 94 °C for 30 s, 52 °C for 90 s, and 68 °C for 2 min, followed by 5 min at 68 °C for one cycle. The primers and primers mixes for multiplex sets are listed at **Table S2**. 5 μl of PCR product was analyzed on a 1% or 2% agarose gel for multiplex PCR for 30 min.

#### Genetic constructions

The genes MMSYN1-0179, MMSYN1-0180 and MMSYN1-0181 were inserted into the plasmid Pmod2-loxpurolox-sp-cre (Hutchison *et al*., 2016) individually and as a cluster together with *mCherry* gene.

For the construction of the plasmid Pmod2-loxpurolox-sp-cre carrying the candidate genes individually, the genes MMSYN1-0179, MMSYN1-0180 and MMSYN1-0181 were amplified with JCVI-syn1.0 genome as the template and the primers pairs Puro_Syn179_fw and Syn179_Puro_rev, Puro_Syn180_fw and Syn180_Puro_rev, Puro_Syn181_fw and Syn181_Puro_rev, named accordingly with their amplicon’s name. The plasmid backbone was amplified using the primers Syn179_Term_fw and Term_Syn179_rev, Syn180_Term_fw and Term_Syn180_rev, Syn181_Term_fw and Term_Syn181_rev, respectively.

The plasmid Pmod2-loxpurolox-sp-cre_mCh-Syn1-0179-0181 was constructed by inserting the genes *mCherry* and the gene cluster MMSYN1-0179-0181 adjacent to the puromycin resistance marker in Pmod2-loxpurolox-sp-cre. The genes MMSYN1-0179-0181 were amplified with JCVI-syn1.0 genome as the template, as well as primers mCh_Syn1-0179_fw and Ter_Syn1-0181_rev. The mCherry gene was amplified from the plasmid pSD024 (Mariscal et al., 2018) and primers puro-mCh_fw and Syn1-0179_mCh_rev. The linear vector Pmod2-loxpurolox-sp-cre was amplified using the plasmid Pmod2-loxpurolox-sp-cre as the template and primers Syn1-0181_Ter_fw and mCh_Puro_rev.

All PCR fragments were amplified using PrimeSTAR Max DNA Polymerase (Takara). The PCR conditions were 98°C 3 min; 30 cycles of 98°C for 10 s, 55°C for 10 s, and 72°C for 2.5 min; and, followed by 72°C for 3 min. PCR products were digested with DpnI (New England BioLabs) to eliminate plasmid template before setting up the assembly reaction using Gibson Assembly ® Master Mix (New England BioLabs). Resulting reaction (2 μL) was transformed by heat shock into NEB 5-alpha Competent *Escherichia coli* (New England BioLabs), following manufacturer’s instructions and plated on LB agar plate with 100 μg/ml ampicillin. The plate was incubated at 37°C overnight. Colony PCR was used to screen positive colonies with genes insertion in the vector using a protocol for OneTaq® Quick-Load® DNA Polymerase (New England BioLabs).

For the insertion of genes MMSYN1-0179 up to MMSYN1-0186 (MMSYN1-0179-0186) into JCVI-syn3B (JCVI-syn3B::mCh-MMSYN1–0179-0186), the plasmid pRS313_loxpurolox-sp-cre_mCh-Syn1-0179-0186 was constructed. The linear vector PRS313 was amplified using the plasmid PRS313 as the template and primers Sp-PRS313_fw and Syn1-0186_PRS313_rev. The genes mCh and MMSYN1-0179-0181 were amplified from the plasmid Pmod2-loxpurolox-sp-cre_mCh-Syn1-0179-0181, using the primers Syn1_182-181_rev and pRS313_Sp_rev. The genes cluster MMSYN1-0182-0183 and MMSYN1-0184-0186 were amplified with JCVI-syn1.0 genome as the template, as well as primers Syn1_181-182_fw and Syn1_184-183_rev, Syn1_183-184_fw and PRS313-Syn186_fw, respectively. The PCR conditions were as previously described. After several unsuccessful attempts to transform *E. coli* with the plasmid assembled in yeast (Gibson, 2011), the DNA fragments and linear vector were assembled via Gibson Assembly®Master Mix (New England BioLabs) and 3 μL of the resulting reaction was used to directly transform JCVI-syn3B.

Primers for PCR amplification, colony PCR and sequencing are listed at **Table S3**. Plasmids maps are shown at **Figure S1**. Sanger sequencing confirmed that the plasmids had no mutations.

#### Transformation of JCVI-syn3B using plasmids and gene expression analysis

Cells were transformed as described previously (Hutchison *et al*., 2016) originating the strains JCVI-syn3B::MMSYN1-0179, JCVI-syn3B::MMSYN1-0180, JCVI-syn3B::MMSYN1-0181, JCVI-syn3B::mCh-MMSYN1–0179-0181 and JCVI-syn3B::mCh-MMSYN1–0179-0186. Briefly, JCVI-syn3B cells were grown in 4 mL of SP4 growth medium to reach pH 6.5 to pH 7.0. The culture was centrifuged for 15 min at 5369 g and 10°C in a 50 mL centrifuge tube. The pellet was resuspended in 3 mL of Sucrose/Tris (S/T) Buffer, composed of 0.5 mol/L sucrose and 10 mmol/L Tris at pH 6.5. The resuspended cells were centrifuged as before. The supernatant was discarded, and the pellet resuspended in 250 μL of 0.1 mol/L CaCl_2_ and incubated for 30 min on ice. Then, 200 ng of plasmid was added to the cells, and the centrifuge tube was mixed gently. 2 mL of 70 % (w/v) polyethylene glycol (PEG) 6000 (Sigma), dissolved in S/T Buffer, was added to the centrifuge tube, and mixed well using a serological pipette. After a 2 min incubation at room temperature, 20 mL S/T Buffer without PEG was added immediately and mixed well. The tube was centrifuged for 15 min at 10,000 g and 8°C. The supernatant was discarded, and the tube inverted with the cap removed on tissue paper to drain residual PEG. The cells were subsequently resuspended in 1 mL of SP4 growth medium prewarmed to 37°C. These cells were incubated for 2 h at 37°C, followed by plating on SP4 agar containing 3 μg/mL of puromycin (Sigma). Colonies appeared after 3 to 4 days at 37°C. Transformations were confirmed by PCR using 1 μL of 1 mL cultures of isolated colonies as template and OneTaq® Quick-Load® DNA Polymerase (New England BioLabs), according with the manufacturer protocol.

#### Live cell fluorescence microscopy

For fluorescence microscopy analysis, we performed an infectivity assay using JCVI-syn3B::mCh-MMSYN1–0179-0181 and JCVI-syn3B::mCh-MMSYN1–0179-0186 add-back mutants, and JCVI-syn1.0 and JCVI-syn3A strains that also expressed mCherry (JCVI-syn1.0::mCh and JCVI-syn3A::mCh, respectively), which allowed us to identify them in co-culture with the mammalian cells. For that, HeLa cells were placed on an 8 well chamber slide at a density of 2 × 10^4^ cells/well and cultured in 500 μl of DMEM (Gibco) with 10% FBS and 100 units/mL of penicillin at 37°C and 5% CO2 for 24 h before use. HeLa cells were initially washed with 500 μl of DPBS and 500 μl of fresh media was added. Cells were inoculated with 5 μl of JCVI-syn1.0::mCh, JCVI-syn3A::mCh, JCVI-syn3B::mCh-MMSYN1–0179-0181, JCVI-syn3B::mCh-MMSYN1–0179-0186, once cultures had entered a stationary growth phase (about 20 h), or 5 μl of DMEM media (no infection).

The cells were incubated at 37°C and 5% CO2 for 24 h or 48h. After incubation cells were treated with 5 μM of CellROX® Green Reagent and incubated at 37°C for 30 min for oxidative stress analysis. The cells were washed 3 times with 500 μl DPBS.

During the last wash 1 drop of NucBlue Live Cell Stain (Hoechst®33342 dye) was added and cells were incubated for 20 min at room temperature (RT) in the dark. After incubation the solution was discarded, and cells were washed once with 500 μl of DPBS. 500 μl of Live Cell Imaging Solution (Life Technologies) was added and the cells were analyzed with the Revolve Fluorescence Microscope (Echo Laboratories Inc., CA, US) at 20x of magnification. All treatments were performed in duplicate.

#### Phagocytosis assay and flow cytometry analysis

A neutrophil infection assay was used to assess the phagocytic activity response induced by the synthetic *Mmc* strains JCVI-syn1.0::mCh, JCVI-syn3A::mCh and the add-back mutants, JCVI-syn3B::mCh-MMSYN1–0179-0181 and JCVI-syn3B::mCh-MMSYN1–0179-0186. Initially, the phagocytic activity of dHL60 in co-culture with JCVI-syn1.0::mCh and JCVI-syn3A::mCh were observed using 3D Cell Explorer Microscope (Nanolive’s 3D Cell Explorer-*fluo*; Model CX-F). 1 ml of synthetic *Mmc* strains broth cultures at the end of exponential phase of growth was spun down at 9000 RCF for 8 min at RT and suspended in 100 μL FBS (Sigma) and used to inoculate 350 μL dHL60 cell suspension (8 × 10^5^ cells/ml) in RPMI media. The co-culture was loaded in an μ-Dish 35 mm (ibidi), and kept at 37°C and 5% CO2 for video acquisition using STEVE software. Images were captured every 1 min for 3 h total duration.

Infected and non-infected neutrophil-like cell were also analyzed by flow cytometry. HL60 control samples and dHL60 cultures were spun down at 275 g for 10 min in RT and suspended in filtered (0.1 μm) RPMI medium at a concentration of 6 × 10^6^ cells/ml. Cell counts were obtained by trypan blue exclusion in the countess cell counter (Invitrogen, CA). 1 ml of HL60 and dHL60 cells were used to suspend synthetic *Mmc* strains prepared as described above and co-cultured at 37°C and 5% CO_2_ up to 210 min. HL60 and dHL60 cells and bacteria alone were used as control samples. Cells were loaded on a BD FACS Aria II (BD Biosciences) equipped with a forward scatter (FCS) photomultiplier tube to obtain raw data in FCS format at 30, 150 and 210 min of co-culture, which were later analyzed by FlowJo v10.6.1 (BD Biosciences). Two samples of each treatment were analyzed over time. To control for extracellular biding of the bacteria to the host cell we place cells on ice and define the quantitative gate. To acquire both bacteria (0.2-0.4 μm diameter) and HL60 and dHL60 cells (~12 μm), FSC and side scatter (SSC) parameters (FSC PMT-A X SSC-A) were set in logarithmic mode and used as threshold signals to collect only bacteria and neutrophils in the scatterplot. mCherry (synthetic *Mmc*) signals were acquired on logarithmic scale, also using the PI-A parameter, and the mean fluorescence intensity (MFI) calculated within a gated region excluding cell debris. A total of 50,000 events were acquired for each sample. Data obtained from less than 50,000 events were not considered in the analysis. Phagocytic activity was expressed as phagocytic index (PI), calculated using the following formula PI = (% phagocytic cells containing ≥ 1 Synthetic *Mmc*) × (mean number of Synthetic *Mmc*/phagocytic cell containing Synthetic *Mmc*) (Barbuddhe et al., 1998). Once host cells were fluorescently labelled post-phagocytosis, MFI from synthetic *Mmc* was used to normalize the PI value in order to allow comparison between samples preserving proportionality.

### Quantification and statistical analysis

Data analyzes were performed using statistical software R (R: A language and environment for statistical computing. R Foundation for Statistical Computing, 2019). As the PI was obtained from the same observational unit at three different times (4 synthetic cell strains × 2 dHL-60 × 3 time points × 2 replicates = 48 observations), the statistical analysis performed was a longitudinal analysis implemented through a linear mixed model (LMM). To perform the analysis, the LMM execution steps for longitudinal data were followed according to Faraway (Faraway, 2016). From the 48 observations, two PI values were not recorded (<50,000 events). Missing data values (JCVI-syn3B::mCh-MMSYN1–0179-0181/DHL60 at 210 min and JCVI-syn3B::mCh-MMSYN1–0179-0186/DHL60 at 30 min) were replaced by similar values of the variable from its matching sample.

We used the Akaike Information Criterion (AIC) (Akaike, 1973) and the Bayesian Information Criterion (BIC) (Schwartz, 1978) for the comparative evaluation among time series models. Four models were evaluated: the first model was the null model Y ~ 1, where Y represents the response variable PI and the number ‘1’ indicates that neither the synthetic cell type nor the incubation time explain the result Y. The second model tested was the LMM with random intercept Y ~ 1 + (1|S), where Y and ‘1’ have the same representation of the null model and (1|S) is the random effect referring to the neutrophil with random intercept. The third model tested was the LMM Y ~ B + (1|S), where Y and (1|S) have the same representation as the previous model, and B represents the fixed effect referring to the type of bacteria. Finally, the fourth model tested was the LMM Y ~ B * T + (1|S) where Y, B and (1|S) have the same representation as the third model and T represents the fixed effect referring to the incubation time. Posteriorly, the significance of fixed effects in the best model identified was evaluated by F tests via Kenward-Roger approximation (Kenward and Roger, 1997). Once the model that satisfactorily represented the sample data was defined, we proceeded with an Honestly Significant Difference test (Tukey, 1949) for multiple comparisons between treatments. For all comparisons, a P-value of <0.05 (95% confidence interval) was considered statistically significant.

## Supplemental Information

**Multimedia files**

**Video S1** - Phagocytic activity of dHL-60 cells in the presence of JCVI-syn1.0::mCh.

**Video S2** - Phagocytic activity of dHL-60 cells in the presence of JCVI-syn3A::mCh.

## Notes

### Competing Interest Statement

The authors have declared no competing interest.

## References

Akaike, H. (1973). Information Theory and an Extension of the Maximum Likelihood Principle. In B.N. Petrov, and Csaki, C., eds. 2nd International Symposium on Information Theory. Akademiai Kiado.

Barbuddhe, S.B., Malik, S.V., and Gupta, L.K. (1998). Effect of in vitro monocyte activation by Listeria monocytogenes antigens on phagocytosis and production of reactive oxygen and nitrogen radicals in bovines. Vet Immunol Immunopathol 64, 149–159. 10.1016/s0165-2427(98)00129-9.

Benders, G.A., Noskov, V.N., Denisova, E.A., Lartigue, C., Gibson, D.G., Assad-Garcia, N., Chuang, R.Y., Carrera, W., Moodie, M., Algire, M.A., et al. (2010). Cloning whole bacterial genomes in yeast. Nucleic acids research 38, 2558–2569. 10.1093/nar/gkq119.

Benedetti, F., Curreli, S., and Zella, D. (2020). Mycoplasmas-Host Interaction: Mechanisms of Inflammation and Association with Cellular Transformation. Microorganisms 8. 10.3390/microorganisms8091351.

Bolske, G. (1988). Survey of Mycoplasma infections in cell cultures and a comparison of detection methods. Zentralbl Bakteriol Mikrobiol Hyg A 269, 331–340. 10.1016/s0176-6724(88)80176-7.

Borchsenius, S.N., Vishnyakov, I.E., Chernova, O.A., Chernov, V.M., and Barlev, N.A. (2020). Effects of Mycoplasmas on the Host Cell Signaling Pathways. Pathogens 9. 10.3390/pathogens9040308.

Breuer, M., Earnest, T.M., Merryman, C., Wise, K.S., Sun, L., Lynott, M.R., Hutchison, C.A., Smith, H.O., Lapek, J.D., Gonzalez, D.J., et al. (2019). Essential metabolism for a minimal cell. Elife 8. 10.7554/eLife.36842.

Browning, G.F., Marenda, M.S., Noormohammadi, A.H., and Markham, P.F. (2011). The central role of lipoproteins in the pathogenesis of mycoplasmoses. Veterinary microbiology 153, 44–50. 10.1016/j.vetmic.2011.05.031.

Cacciotto, C., Cubeddu, T., Addis, M.F., Anfossi, A.G., Tedde, V., Tore, G., Carta, T., Rocca, S., Chessa, B., Pittau, M., and Alberti, A. (2016). Mycoplasma lipoproteins are major determinants of neutrophil extracellular trap formation. Cell Microbiol 18, 1751–1762. 10.1111/cmi.12613.

Cherry, J.D., and Taylor-Robinson, D. (1970). Growth and Pathogenesis of Mycoplasma mycoides var. capri in Chicken Embryo Tracheal Organ Cultures. Infection and immunity 2, 431–438. 10.1128/iai.2.4.431-438.1970.

Collins, S.J., Ruscetti, F.W., Gallagher, R.E., and Gallo, R.C. (1978). Terminal differentiation of human promyelocytic leukemia cells induced by dimethyl sulfoxide and other polar compounds. Proceedings of the National Academy of Sciences of the United States of America 75, 2458–2462. 10.1073/pnas.75.5.2458.

Davies, J.S., Currie, M.J., Wright, J.D., Newton-Vesty, M.C., North, R.A., Mace, P.D., Allison, J.R., and Dobson, R.C.J. (2021). Selective Nutrient Transport in Bacteria: Multicomponent Transporter Systems Reign Supreme. Front Mol Biosci 8, 699222. 10.3389/fmolb.2021.699222.

Dedieu, L., and Balcer-Rodrigues, V. (2006). Viable Mycoplasma mycoides ssp. mycoides small colony-mediated depression of the bovine cell responsiveness to the mitogen concanavalin A. Scandinavian journal of immunology 64, 376–381. 10.1111/j.1365-3083.2006.01799.x.

Di Teodoro, G., Marruchella, G., Di Provvido, A., D’Angelo, A.R., Orsini, G., Di Giuseppe, P., Sacchini, F., and Scacchia, M. (2020). Contagious Bovine Pleuropneumonia: A Comprehensive Overview. Vet Pathol 57, 476–489. 10.1177/0300985820921818.

Di Teodoro, G., Marruchella, G., Di Provvido, A., Orsini, G., Ronchi, G.F., D’Angelo, A.R., D’Alterio, N., Sacchini, F., and Scacchia, M. (2018). Respiratory explants as a model to investigate early events of contagious bovine pleuropneumonia infection. Vet Res 49, 5. 10.1186/s13567-017-0500-z.

Faraway, J.J. (2016). Extending the Linear Model with R - Generalized Linear, Mixed Effects and Nonparametric Regression Models, 2nd Edition Edition (CRC Press).

Fraser, C.M., Gocayne, J.D., White, O., Adams, M.D., Clayton, R.A., Fleischmann, R.D., Bult, C.J., Kerlavage, A.R., Sutton, G., Kelley, J.M., et al. (1995). The minimal gene complement of Mycoplasma genitalium. Science 270, 397–403.

Gibson, D.G. (2011). Gene and genome construction in yeast. Curr Protoc Mol Biol Chapter 3, Unit3 22. 10.1002/0471142727.mb0322s94.

Gibson, D.G., Glass, J.I., Lartigue, C., Noskov, V.N., Chuang, R.Y., Algire, M.A., Benders, G.A., Montague, M.G., Ma, L., Moodie, M.M., et al. (2010). Creation of a bacterial cell controlled by a chemically synthesized genome. Science 329, 52–56. 10.1126/science.1190719.

Hernandez, L., Lopez, J., St-Jacques, M., Ontiveros, L., Acosta, J., and Handel, K. (2006). Mycoplasma mycoides subsp. capri associated with goat respiratory disease and high flock mortality. Can Vet J 47, 366–369.

Hossain, T., Deter, H.S., Peters, E.J., and Butzin, N.C. (2021). Antibiotic tolerance, persistence, and resistance of the evolved minimal cell, Mycoplasma mycoides JCVI-syn3B. iScience 24, 102391. 10.1016/j.isci.2021.102391.

Hutchison, C.A. 3rd, Chuang, R.Y., Noskov, V.N., Assad-Garcia, N., Deerinck, T.J., Ellisman, M.H., Gill, J., Kannan, K., Karas, B.J., Ma, L., et al. (2016). Design and synthesis of a minimal bacterial genome. Science 351, aad6253. 10.1126/science.aad6253.

Ji, Y., Karbaschi, M., and Cooke, M.S. (2019). Mycoplasma infection of cultured cells induces oxidative stress and attenuates cellular base excision repair activity. Mutat Res Genet Toxicol Environ Mutagen 845, 403054. 10.1016/j.mrgentox.2019.05.010.

Jores, J., Ma, L., Ssajjakambwe, P., Schieck, E., Liljander, A., Chandran, S., Stoffel, M.H., Cippa, V., Arfi, Y., Assad-Garcia, N., et al. (2019). Removal of a Subset of Non-essential Genes Fully Attenuates a Highly Virulent Mycoplasma Strain. Frontiers in microbiology 10, 664. 10.3389/fmicb.2019.00664.

Kenward, M.G., and Roger, J.H. (1997). Small sample inference for fixed effects from restricted maximum likelihood. Biometrics 53, 983–997.

Kong, F., James, G., Gordon, S., Zelynski, A., and Gilbert, G.L. (2001). Species-specific PCR for identification of common contaminant mollicutes in cell culture. Applied and environmental microbiology 67, 3195–3200. 10.1128/AEM.67.7.3195-3200.2001.

Lartigue, C., Vashee, S., Algire, M.A., Chuang, R.Y., Benders, G.A., Ma, L., Noskov, V.N., Denisova, E.A., Gibson, D.G., Assad-Garcia, N., et al. (2009). Creating bacterial strains from genomes that have been cloned and engineered in yeast. Science 325, 1693–1696. 10.1126/science.1173759.

Manfredi, A.A., Ramirez, G.A., Rovere-Querini, P., and Maugeri, N. (2018). The Neutrophil’s Choice: Phagocytose vs Make Neutrophil Extracellular Traps. Front Immunol 9, 288. 10.3389/fimmu.2018.00288.

Mariscal, A.M., Kakizawa, S., Hsu, J.Y., Tanaka, K., Gonzalez-Gonzalez, L., Broto, A., Querol, E., Lluch-Senar, M., Pinero-Lambea, C., Sun, L., et al. (2018). Tuning Gene Activity by Inducible and Targeted Regulation of Gene Expression in Minimal Bacterial Cells. ACS synthetic biology. 10.1021/acssynbio.8b00028.

Medzhitov, R. (2007). Recognition of microorganisms and activation of the immune response. Nature 449, 819–826. 10.1038/nature06246.

Morowitz, H.J. (1984). The completeness of molecular biology. Israel journal of medical sciences 20, 750–753.

Nguyen, M.T., Matsuo, M., Niemann, S., Herrmann, M., and Gotz, F. (2020). Lipoproteins in Gram-Positive Bacteria: Abundance, Function, Fitness. Frontiers in microbiology 11, 582582. 10.3389/fmicb.2020.582582.

Nicholas, R.A., Ayling, R.D., and McAuliffe, L. (2009). Vaccines for Mycoplasma diseases in animals and man. J Comp Pathol 140, 85–96. 10.1016/j.jcpa.2008.08.004.

Nikfarjam, L., and Farzaneh, P. (2012). Prevention and detection of Mycoplasma contamination in cell culture. Cell J 13, 203–212.

Nishiumi, F., Kawai, Y., Nakura, Y., Yoshimura, M., Wu, H.N., Hamaguchi, M., Kakizawa, S., Suzuki, Y., Glass, J.I., and Yanagihara, I. (2021). Blockade of endoplasmic reticulum stress-induced cell death by Ureaplasma parvum vacuolating factor. Cell Microbiol, e13392. 10.1111/cmi.13392.

Noccard, E., and Roux, P. (1898). Le microbe de la peripneumonie. Annales de l’Institut Pasteur 12, 240–262.

Pelletier, J.F., Sun, L., Wise, K.S., Assad-Garcia, N., Karas, B.J., Deerinck, T.J., Ellisman, M.H., Mershin, A., Gershenfeld, N., Chuang, R.Y., et al. (2021). Genetic requirements for cell division in a genomically minimal cell. Cell 184, 2430–2440 e2416. 10.1016/j.cell.2021.03.008.

Pilo, P., Frey, J., and Vilei, E.M. (2007). Molecular mechanisms of pathogenicity of Mycoplasma mycoides subsp. mycoides SC. Vet J 174, 513–521. 10.1016/j.tvjl.2006.10.016.

R: A language and environment for statistical computing. R Foundation for Statistical Computing (2019). (R Core Team).

Razin, S., Yogev, D., and Naot, Y. (1998). Molecular biology and pathogenicity of mycoplasmas. Microbiology and molecular biology reviews : MMBR 62, 1094–1156. 10.1128/MMBR.62.4.1094-1156.1998.

Rosales, C. (2020). Neutrophils at the crossroads of innate and adaptive immunity. J Leukoc Biol 108, 377–396. 10.1002/JLB.4MIR0220-574RR.

Rottem, S. (2003). Interaction of mycoplasmas with host cells. Physiol Rev 83, 417–432. 10.1152/physrev.00030.2002.

Sasaki, Y., Ishikawa, J., Yamashita, A., Oshima, K., Kenri, T., Furuya, K., Yoshino, C., Horino, A., Shiba, T., Sasaki, T., and Hattori, M. (2002). The complete genomic sequence of Mycoplasma penetrans, an intracellular bacterial pathogen in humans. Nucleic acids research 30, 5293–5300.

Sauer, B. (1987). Functional expression of the cre-lox site-specific recombination system in the yeast Saccharomyces cerevisiae. Mol Cell Biol 7, 2087–2096. 10.1128/mcb.7.6.2087-2096.1987.

Schwartz, G. (1978). Estimating the Dimension of a Model. The Anals of Statistics 6, 461–464.

Scieszka, D., Lin, Y.-H., Li, W., Choudhury, S., Yu, Y., and Freire, M. (2020). Netome: The Molecular Characterization of Neutrophil Extracellular Traps (NETs). bioRxiv (preprint). https://doi.org/10.1101/2020.05.18.102772.

Shimizu, T. (2016). Inflammation-inducing Factors of Mycoplasma pneumoniae. Frontiers in microbiology 7, 414. 10.3389/fmicb.2016.00414.

Steiner, T., and McGarrity, G. (1983). Mycoplasmal infection of insect cell cultures. In Vitro 19, 672–682. 10.1007/BF02628958.

Tambi, N.E., Maina, W.O., and Ndi, C. (2006). An estimation of the economic impact of contagious bovine pleuropneumonia in Africa. Rev Sci Tech 25, 999–1011.

Tan, M.F., Gao, T., Liu, W.Q., Zhang, C.Y., Yang, X., Zhu, J.W., Teng, M.Y., Li, L., and Zhou, R. (2015). MsmK, an ATPase, Contributes to Utilization of Multiple Carbohydrates and Host Colonization of Streptococcus suis. PLoS One 10, e0130792. 10.1371/journal.pone.0130792.

Tang, J., Hu, M., Lee, S., and Roblin, R. (2000). A polymerase chain reaction based method for detecting Mycoplasma/Acholeplasma contaminants in cell culture. J Microbiol Methods 39, 121–126. 10.1016/s0167-7012(99)00107-4.

Thiam, H.R., Wong, S.L., Qiu, R., Kittisopikul, M., Vahabikashi, A., Goldman, A.E., Goldman, R.D., Wagner, D.D., and Waterman, C.M. (2020). NETosis proceeds by cytoskeleton and endomembrane disassembly and PAD4-mediated chromatin decondensation and nuclear envelope rupture. Proceedings of the National Academy of Sciences of the United States of America 117, 7326–7337. 10.1073/pnas.1909546117.

Thiaucourt, F., and Bolske, G. (1996). Contagious caprine pleuropneumonia and other pulmonary mycoplasmoses of sheep and goats. Rev Sci Tech 15, 1397–1414. 10.20506/rst.15.4.990.

Timenetsky, J., Santos, L.M., Buzinhani, M., and Mettifogo, E. (2006). Detection of multiple mycoplasma infection in cell cultures by PCR. Braz J Med Biol Res 39, 907–914. 10.1590/s0100-879x2006000700009.

Tseng, C.W., Kanci, A., Citti, C., Rosengarten, R., Chiu, C.J., Chen, Z.H., Geary, S.J., Browning, G.F., and Markham, P.F. (2013). MalF is essential for persistence of Mycoplasma gallisepticum in vivo. Microbiology (Reading) 159, 1459–1470. 10.1099/mic.0.067553-0.

Tukey, J.W. (1949). Comparing individual means in the analysis of variance. Biometrics 5, 99–114.

Williamson, D.L., and Whitcomb, R.F. (1975). Plant mycoplasmas: a cultivable spiroplasma causes corn stunt disease. Science 188, 1018–1020.

Woese, C.R., Maniloff, J., and Zablen, L.B. (1980). Phylogenetic analysis of the mycoplasmas. Proceedings of the National Academy of Sciences of the United States of America 77, 494–498. 10.1073/pnas.77.1.494.

Yiwen, C., Yueyue, W., Lianmei, Q., Cuiming, Z., and Xiaoxing, Y. (2021). Infection strategies of mycoplasmas: Unraveling the panoply of virulence factors. Virulence 12, 788–817. 10.1080/21505594.2021.1889813.

