## Supplementary Material for "Minimal cell JCVI-syn3B as a chassis to reveal the mechanisms behind *Mycoplasma* host–pathogen interactions"

**Table S1 -** Genome characteristics of synthetic *Mycoplasma* strains used in this work, *related to STAR methods*.

| ***Mycoplasma* strains** | **Genome size (kb)** | **Total genes** | ***MMSYN1-179-186* are present in the genome?** |
|---|---|---|---|
| *M. mycoides* JCVI-Syn1.0[a] | 1080 | 901 | Yes |
| *M. mycoides* JCVI-Syn2.0[b] | 576 | 516 | No |
| *M. mycoides* JCVI-Syn3.0[c] | 531 | 473 | No |
| *M. mycoides* JCVI-Syn3A[d] | 543 | 491 | No |
| *M. mycoides* JCVI-Syn3B[e] | 544 | 491 | No |
| *M. mycoides* JCVI-Syn1.0F[f] | 1067 | 893 | No |
| RDG1.0-1[c] | 999 | 876 | Yes |
| RDG1.0-2[c] | 1029 | 862 | No |
| RDG1.0-3[c] | 1006 | 877 | Yes |
| RDG1.0-4[c] | 997 | 885 | Yes |
| RDG1.0-5[c] | 1023 | 878 | Yes |
| RDG1.0-6[c] | 999 | 873 | Yes |
| RDG1.0-7[c] | 1025 | 862 | Yes |
| RDG1.0-8[c] | 1013 | 873 | Yes |
| RGD1.0-2,3,5,6,7,8 + syn1.0-1,4 | 740 | 567 | No |

[a]Gibson et al. 2010. (GenBank CP002027.1)

[b]Hutchison et al. 2016. (GenBank CP014992.1)

[c]Hutchison et al. 2016. (GenBank CP014940.1)

[d]Breuer et al., 2019. (GenBank CP016816.2)

[e]Nishiumi et al., 2021 (GenBank accession number being obtained)

**Table S2 –** List of primers and primers mixes for multiplex PCR used for Color Changing Unit (CCU) validation, *related to STAR methods*.

| Primer name | Primer sequence |
|---|---|
| JCVI-syn 23S rRNA_fw | AGCAGTCAGAACATGGGTGA |
| JCVI-syn 23S rRNA_rev | TAGCTGTCTGTGCAGTTCCA |
| SeqSyn180a_fw | GCTGCTGATAAAGTCAATAAAAG |
| SeqSyn180b_rev | CATAACTGTTGTTCCATCTC |
| RCO1776 | ACCAAGTACAATGCTAAGTG |
| RCO1777 | CTCCTGAATACTTATTTGAAAAAC |
| SBIR-2F | TTTTAGTATGTTTAAGGAGAAGA |
| MPCR9-2R | TGATGGTGCATAACGTAATC |
| MPCR11-3F | TGATGATGATAATAAATTAAATCCT |
| RCO1778 | AAAAAGCTATTTTTTACAAGTTCAA |
| RCO1779 | TATTAGGATCAGTAGCTAAAGG |
| RCO1780 | GTTTAAGTCAATCTTTTCTTCTTG |
| RCO1781 | ACCTATTATTTATAAATAACAATGC |
| RCO1782 | TTGTAGCAACTGGATATAATGG |
| SBIR-6F | GTTTTACCAACTCCAGGTTC |
| MPCR9-6R | TGCAAAGACTAAAGTATATGTG |
| RCO1805N | AACTCCTATGGGTGTGTTG |
| RCO1784 | TAAATTATTATGACAATTCACTTTC |
| RCO1785 | GACTATTGCCTCCAATTAGA |
| RCO1786 | TCTTTTGCTAATTGTTGACTTG |

**Table S3 -** List of primers used for plasmids construction and evaluation (Seq primers; named according with its target), *related to STAR methods*.

| Primer name | Primer sequence |
|---|---|
| Puro_Syn179_Fw | CTTGGTGTATGACTAGAAAACCTGGTGCTTAACCATCATTTCTCCTTTTT |
| Syn179_Puro_Rev | AAAAAGGAGAAATGATGGTTAAGCACCAGGTTTTCTAGTCATACACCAAG |
| Syn179_Term_Fw | CACAATATAAGTTCCATTAACAAAAAAATCGGGAAATCCCG |
| Term_Syn179_Rev | CGGGATTTCCCGATTTTTTTGTTAATGGAACTTATATTGTG |
| Puro_Syn180_Fw | CTTGGTGTATGACTAGAAAACCTGGTGCTTAATAAAAAGAATTAAAAAC |
| Syn180_Puro_Rev | GTTTTTAATTCTTTTTATTAAGCACCAGGTTTTCTAGTCATACACCAAG |
| Syn180_Term_Fw | CTGAAAAATTAAAATACAAATTCCACTAACAAAAAAATCGGGAAATC |
| Term_Syn180_Rev | GATTTCCCGATTTTTTTGTTAGTGGAATTTGTATTTTAATTTTTCAG |
| Puro_Syn181_Fw | GACTAGAAAACCTGGTGCTTAAATGGACTTTCAACAAGCTTTTG |
| Syn181_Puro_Rev | CAAAAGCTTGTTGAAAGTCCATTTAAGCACCAGGTTTTCTAGTC |
| Syn181_Term_Fw | GAGAATATAGATTTTAGAAAGTGAGGCAAAAAAATCGGGAAATC |
| Term_Syn181_Rev | GATTTCCCGATTTTTTTGCCTCACTTTCTAAAATCTATATTCTC |
| mCh_Syn1-179_fw | CTACTGGTGGTATGGATGAACTATATAAATAACCATCATTTCTCCTTTTT |
| Ter_Syn1-181_rev | CGG GAT TTC CCG ATT TTT TTG TAA CCT CAC TTT CTA AAA TCT ATA TTC TC |
| Puro-mCh_fw | CTTGGTGTATGACTAGAAAACCTGGTGCTTAAATGGCTATTATTAAAGAATTTATGAG |
| Syn1-179_mCh_rev | AAAAAGGAGAAATGATGGTTATTTATATAGTTCATCCATACCACCAGTAG |
| Syn1-181_Ter_fw | GAG AAT ATA GAT TTT AGA AAG TGA GGT TAC AAA AAA ATC GGG AAA TCC CG |
| mCh_Puro_rev | CTCATAAATTCTTTAATAATAGCCATTTAAGCACCAGGTTTTCTAGTCATACACCAAG |
| Sp-PRS313_fw | CAGGGTTGAGATGTGTATAAGGCAAGGCGATTAAGTTGGG |
| Syn1-186_PRS313_rev | GAACACCCGAGAAAATTCATACATCCCCCCTTCGCCAGC |
| Syn1_182-181_rev | CTCTATATGATTCATATTAACCTCACTTTCTAAAATCTATATTCTCTTTTTTTCATTTTT |
| pRS313_Sp_rev | CCCAACTTAATCGCCTTGCCTTATACACATCTCAACCCTG |
| Syn1_181-182_fw | AAAAATGAAAAAAAGAGAATATAGATTTTAGAAAGTGAGGTTAATATGAATCATAT AGAG |
| Syn1_184-183_rev | CTTAATTCAACTTTCATACTAACCTTTTACTCCTCCAGATAATCCACCTG |
| Syn1_183-184_fw | CAGGTGGATTATCTGGAGGAGTAAAAGGTTAGTATGAAAGTTGAATTAAG |
| PRS313-Syn186_fw | GCTGGCGAAGGGGGGATGTATGAATTTTCTCGGGTGTTC |
| SeqPuro_fw | GAAGTTCCTGAAGGTCCTAG |
| SeqSyn179_fw | CAT TTG TTG ACG CTG GTG |
| SeqSyn179_rev | CAC CAG CGT CAA CAA ATG |
| SeqSyn1_180_fw | CAGTTATGTCGCTTGTTGG |
| SeqSyn180a_fw | GCTGCTGATAAAGTCAATAAAAG |
| SeqSyn180a_rev | CTTTTATTGACTTTATCAGCAGC |
| SeqSyn180b_fw | GAGATGGAACAACAGTTATG |
| SeqSyn180b_rev | CATAACTGTTGTTCCATCTC |
| SeqSyn181_fw | CCAAGCATAATCAGTATTG |
| SeqSyn181_rev | CAATACTGATTATGCTTGG |
| SeqSyn1_182_fw | GGTACTTCAGCTAATGGAT |
| SeqSyn182_fw | GAGCAGATAATCCTGATGG |

| | |
|---|---|
| SeqSyn182_rev | CCATCAGGATTATCTGCTC |
| SeqSyn183a_fw | CTGATTCTAAGCTAATTGAAGG |
| SeqSyn183a_rev | CCTTCAATTAGCTTAGAATCAG |
| SeqSyn183b_fw | CAGATAAACCACCACTAAG |
| SeqSyn183b_rev | CTTAGTGGTGGTTTATCTG |
| SeqSyn184_fw | CTGGAGGTCAACAACAAC |
| SeqSyn184_rev | GTTGTTGTTGACCTCCAG |
| SeqSyn185a_fw | GGATGCTATGTTGGTGG |
| SeqSyn185a_rev | CCACCAACATAGCATCC |
| SeqSyn185b_fw | GGAGTTACTGGTCCTAATG |
| SeqSyn185b_rev | CATTAGGACCAGTAACTCC |
| SeqSyn1_186_fw | TGAAGTGATGAGCAAGTAG |
| SeqSyn186_fw | GCAAAAGAACAGCCTGA |
| SeqSyn186_rev | TCAGGCTGTTCTTTTGC |
| SeqTerm_rev | GGTGTTCTCGCATATTG |
| SeqmCh_fw | GGAGGACATTATGATGCAG |
| SeqmCherry_rev | GTCTTCCTTCTCCTTCAC |
| Seq-Ori_rev | CAATGCTCACGCTGTAGGTATC |
| Seq-pRS313_rev | CTCACTAAAGGGAACAAAAGC |
| Seq-f1ori_rev | GAGTGTTGTTCCAGTTTGG |

**Table S4** – Values of Akaike Information Criterion (AIC) and the Bayesian Information Criterion (BIC) for the comparative evaluation among time series models representing phagocytic index (Y) in function of synthetic cell type (B) and time of analysis (T), *related to Quantification and statistical analysis*. Number '1' indicates that neither the synthetic cell type nor the incubation time explain the result Y and (1|S) is the random effect referring to the neutrophil with random intercept.

| | Model | AIC | BIC |
|---|---|---|---|
| (1) | Y ~ 1 | 291,6 | 295,2 |
| (2) | Y ~ 1 + (1\|S) | 247,9 | 253,4 |
| (3) | Y ~ B + (1\|S) | 221,4 | 239,7 |
| (4) | Y ~ B * T + (1\|S) | 187,2 | 234,7 |
| (5) | Y ~ B * T + (T\|S) | - | - |

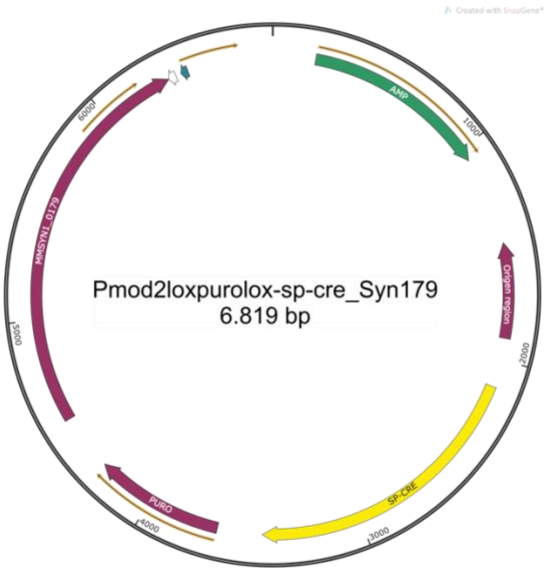
Pmod2loxpurolox-sp-cre_Syn179
6.819 bp
AMP
1000
Origen region
2000
SP-CRE
3000
PURO
4000
5000
MMSYN1_0179
6000

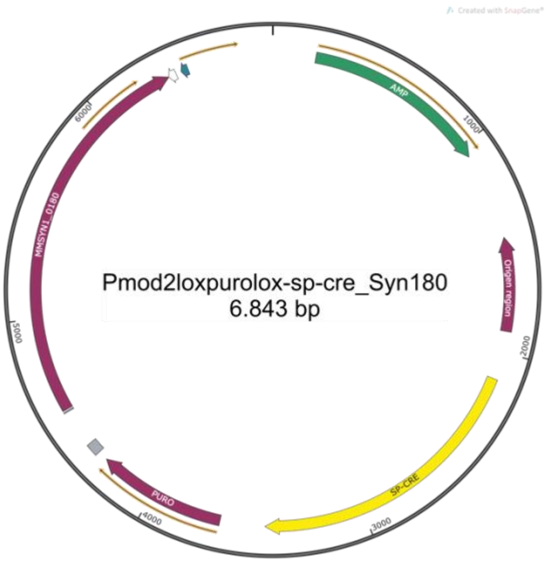
Pmod2loxpurolox-sp-cre_Syn180
6.843 bp
AMP
1000
Origen region
2000
SP-CRE
3000
PURO
4000
5000
MMSYN1_0180
6000

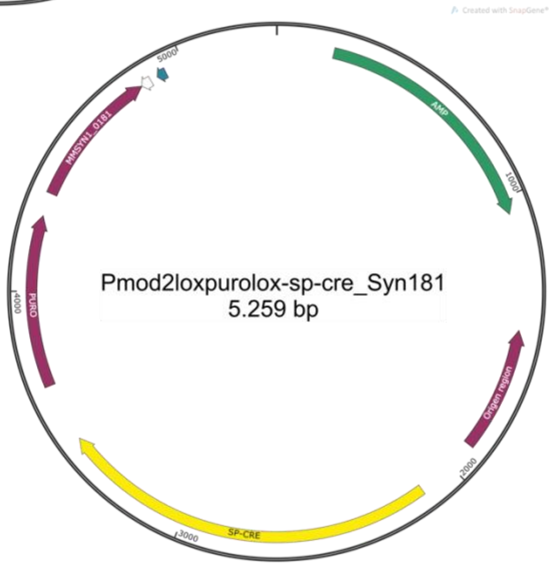
Pmod2loxpurolox-sp-cre_Syn181
5.259 bp
AMP
1000
Origen region
2000
SP-CRE
3000
PURO
4000
MMSYN1_0181
5000

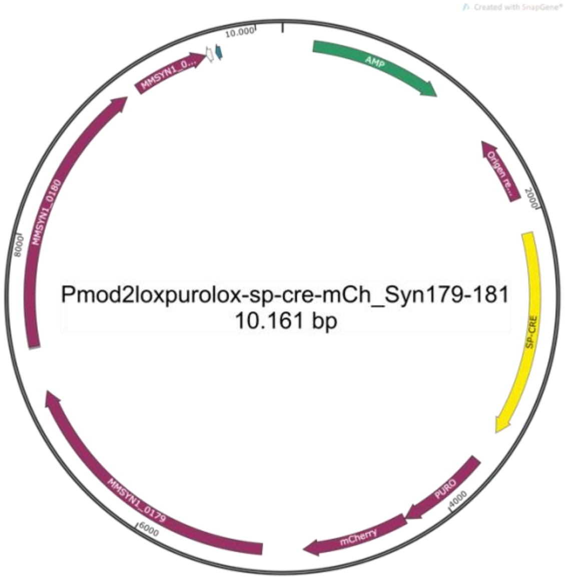
Pmod2loxpurolox-sp-cre-mCh_Syn179-181
10.161 bp
AMP
Origen re...
2000
SP-CRE
PURO
4000
mCherry
MMSYN1_0179
6000
8000
MMSYN1_0180
MMSYN1_0...
10.000

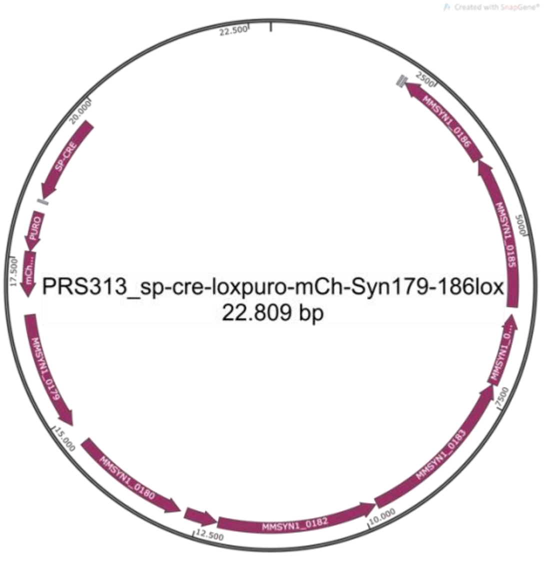
PRS313_sp-cre-loxpuro-mCh-Syn179-186lox
22.809 bp
22.500
2500
MMSYN1_0186
5000
MMSYN1_0185
MMSYN1_0...
7500
MMSYN1_0183
10.000
MMSYN1_0182
12.500
MMSYN1_0180
15.000
MMSYN1_0179
17.500
mCh...
PURO
SP-CRE
20.000

**Figure S1 – Plasmids constructed in this study,** *related to STAR methods*. Maps created using SnapGene® software (from Insightful Science; available at snapgene.com).

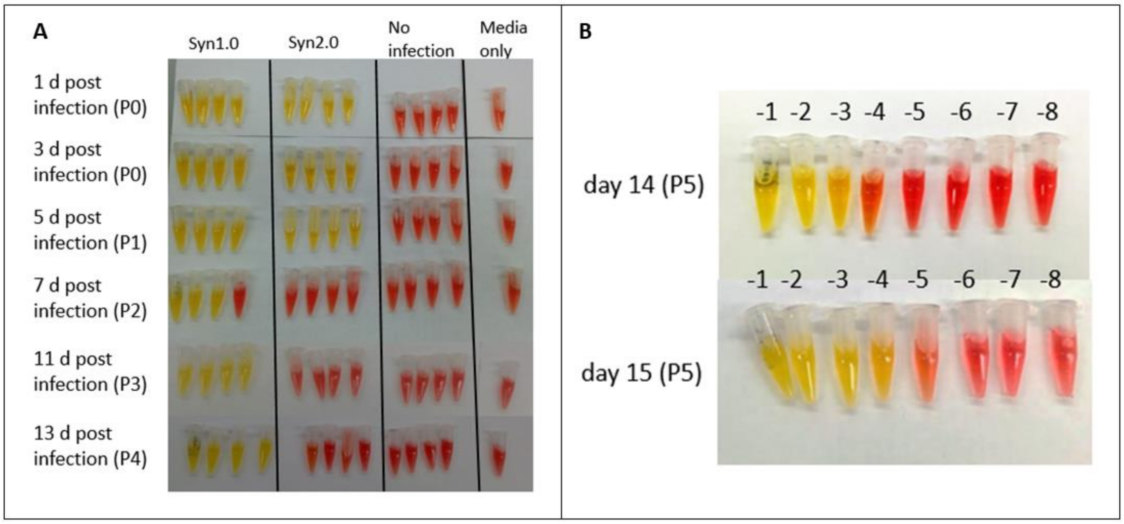


**Figure S2 - CCU assay of synthetic *Mmc* infection in HEK-293T cells,** *related to Detection of synthetic Mmc cell in co-culture with mammalian cells*. (A) 100 µl of an actively growing culture of JCVI-syn 1.0 (Syn 1.0), JCVI-syn 2.0 (Syn 2.0), or 100 µl of DMEM media (No infection) were added to a culture of HEK-293T cells. At selected times over 13 days a sample of supernatant was removed and a CCU assay was performed. Uninfected media is shown as a negative control to show the color without any bacterial growth. For each time point four 10X serial dilutions were performed. The lowest dilutions are on the left and most diluted on the right. Passage numbers (indicating the number of times the HEK-293T cells had been passaged since mixed with mycoplasmas) is given in parentheses. (B) JCVI-syn1.0 grows with HEK-293T cells. A CCU assay was performed on day 14 and day 15 after a new passage (P5). The increase in mycoplasma titer between 14 and 15 days indicated the JCVI-syn1.0 was growing. In all instances, images of the CCU assays were taken after 24 h of incubation. $P_0$ refers to HEK-293T cell culture passage zero.

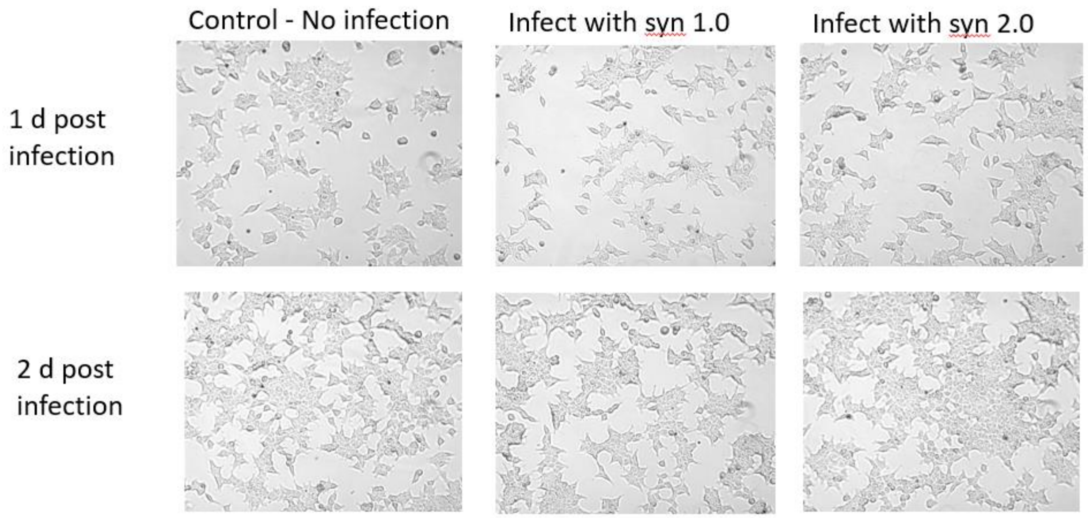


**Figure S3 - Synthetic *M. mycoides* growth does not affect morphology in HEK-293T cells,** *related to Detection of synthetic Mmc cell in co-culture with mammalian cells*. Micrograph images of HEK-293T were taken using an inverted scope at 30x magnification at one and two days of co-culture with JCVI-syn1.0 and JCVI-syn2.0.

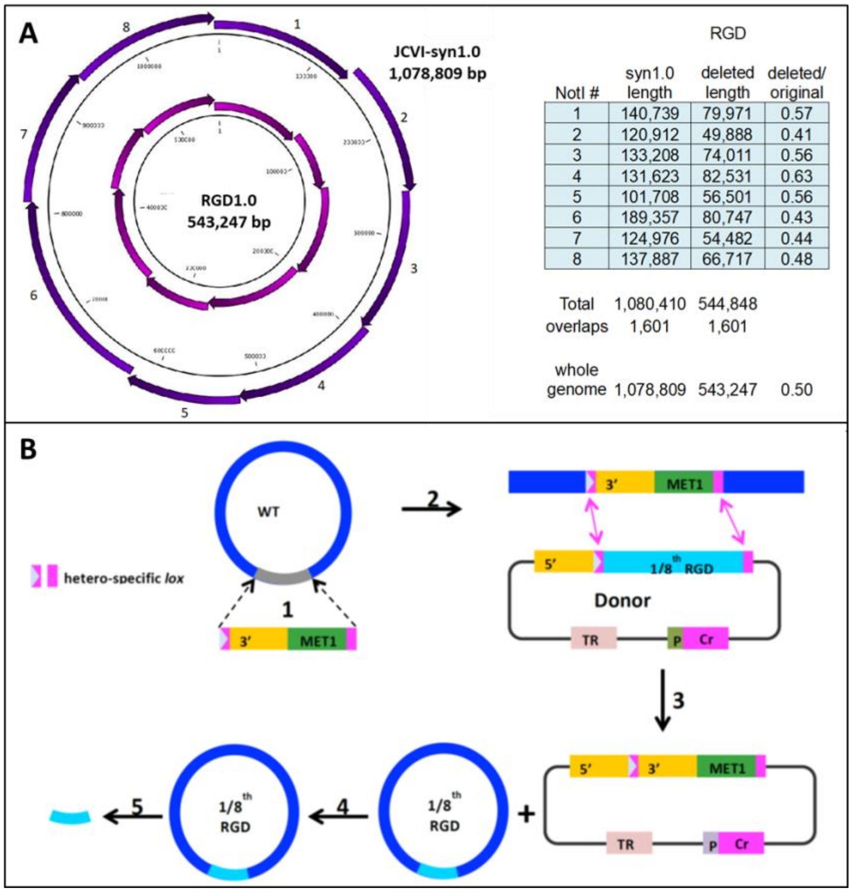


**Figure S4 – Method for construction of 8 different genomes in which only 1/8th of the genome is reduced and 7/8ths are wild type,** related to *Using minimized Mmc strains to identify candidate genes necessary for infection*. The Reduced Genome Design (RGD) set, called RGD1.0, is comprised of 8 strains whose genomes consist of a 1/8th RGD segment and 7/8ths syn1.0 genome were constructed and transplanted (A). A swapping approach, based on recombinase-mediated cassette exchange (RMCE), was developed to improve the efficiency of genome construction (18). An approximate 1/8th genome (colored with gray) of JCVI-syn1.0 cloned in yeast was replaced with a landing pad cassette flanked by two 34-bp hetero-specific *loxP* sites. (B) (2) A donor plasmid containing a corresponding 1/8th RGD, flanked by another two hetero-specific lox sites was introduced into to the landing pad strain by yeast mating. 3) The 1/8th RGD segment was exchanged with the landing pad locus via Cremediated recombination. An intron containing a URA3 gene was split between the landing pad and the donor plasmid so that the cassette exchange could be selected by restoration of uracil prototrophy. 4) The $1/8^{th}$ RGD + $7/8^{ths}$ JCVI-syn1.0 genome was transplanted into *M. capricolum* recipient cells to produce a *M. mycoides* strain. 5) A $1/8^{th}$ RGD fragment can be purified by NotI digestion and then used for genome assembly. Figure from Hutchison *et al*., 2016.

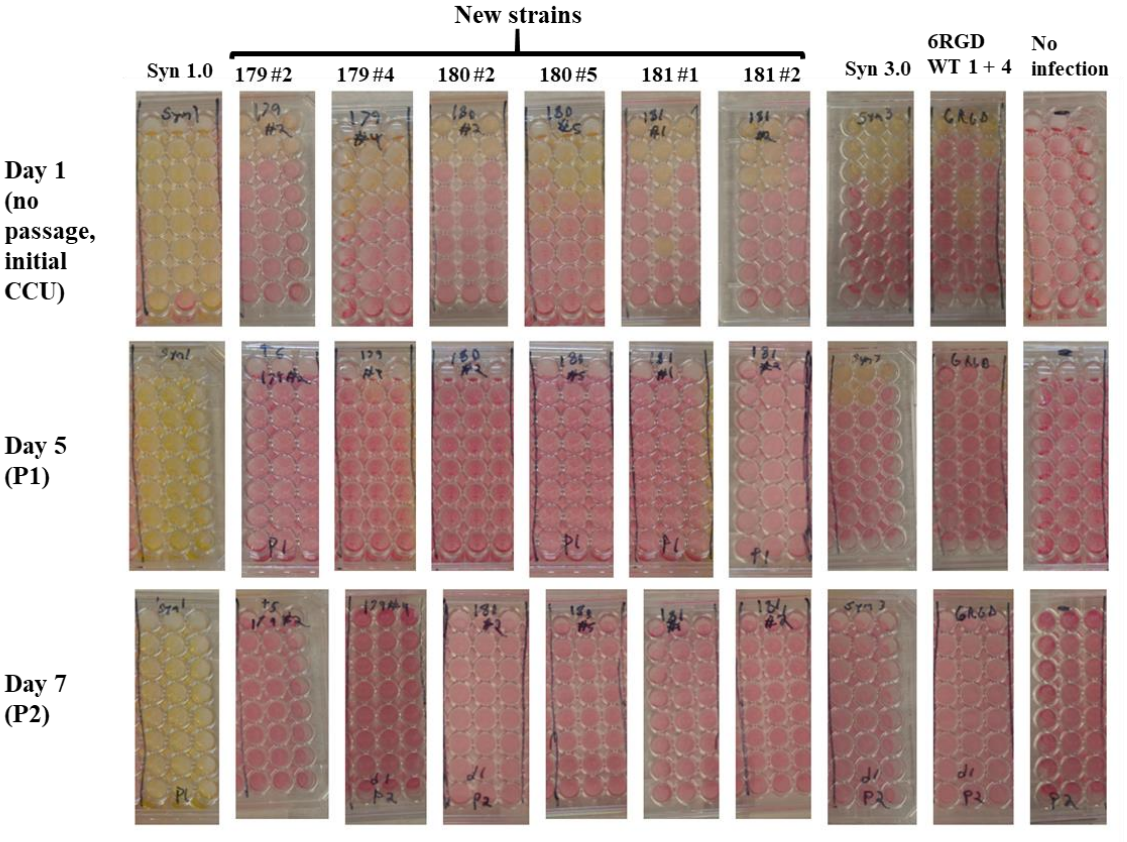


**Figure S5 - CCU assay of synthetic *Mmc* JCVI-syn3B add-back mutants mixed with in HEK-293T cells**, related to *Construction of JCVI-syn3B add-back mutants and CCU analysis after their incubation with mammalian cells.* No mycoplasma survival was observed from the New strains JCVI-syn3B::MMSYN1-179 (179), JCVI-syn3B::MMSYN1-180 (180) and JCVI-syn3B::MMSYN1-181 (181) are JCVI-syn3B with either MMSYN1-179, MMSYN1-180, or MMSYN1-181 inserted into the dual *loxP* landing pad so that the genes were expressed. Two clones of each of those strains were analyzed by mixing separately with HEK-293T cells. The same mixing with HEK-293T cells procedures was done with JCVI-SYN1.0 (Syn1.0), JCVI-syn3.0 (Syn 3.0) and a strain containing six RGD1.0 1/8$^{th}$ genome segments and JCVI-syn1.0 1/8$^{th}$ genome segments 1 and 4 (6 RGD WT 1 + 4). The mixtures of HEK-293T cells and mycoplasma strains were incubated at 37°C and 5% $CO_2$ for 10 days and then CCU assays were performed. The CCU assays were incubated at 37°C for 7 days and then read. The growth media in the JCVI-syn1.0 (Syn 1.0) HEK-293T cells mixture contained more than $10^{10}$ bacteria per ml. None of the other mycoplasma strains survived the 10 day incubation with HEK-293T cells as indicated by the red color of the SP4 media in all the CCU wells. The “No infection” samples are CCUs from HEK-293T cells culture mixed with DMEM media. For the CCU assay, eight 10X serial dilutions were performed (the tops of the 96 well plates contain the $10^{-1}$ dilution and the next rows down are progressively greater dilutions). As evidenced by the three 8 well columns shown for each mycoplasma sample, the CCU determinations were done in triplicate.

**Figure S6 - Gating strategy for flow cytometry data analysis,** *related to Figures 5 and 6.* (A, B, C and D) Representative flow cytometry plots are for synthetic mycoplasmas in the presence of dHL-60 and HL-60 cells after 30 min, 90 min and 210 min of co-culture.

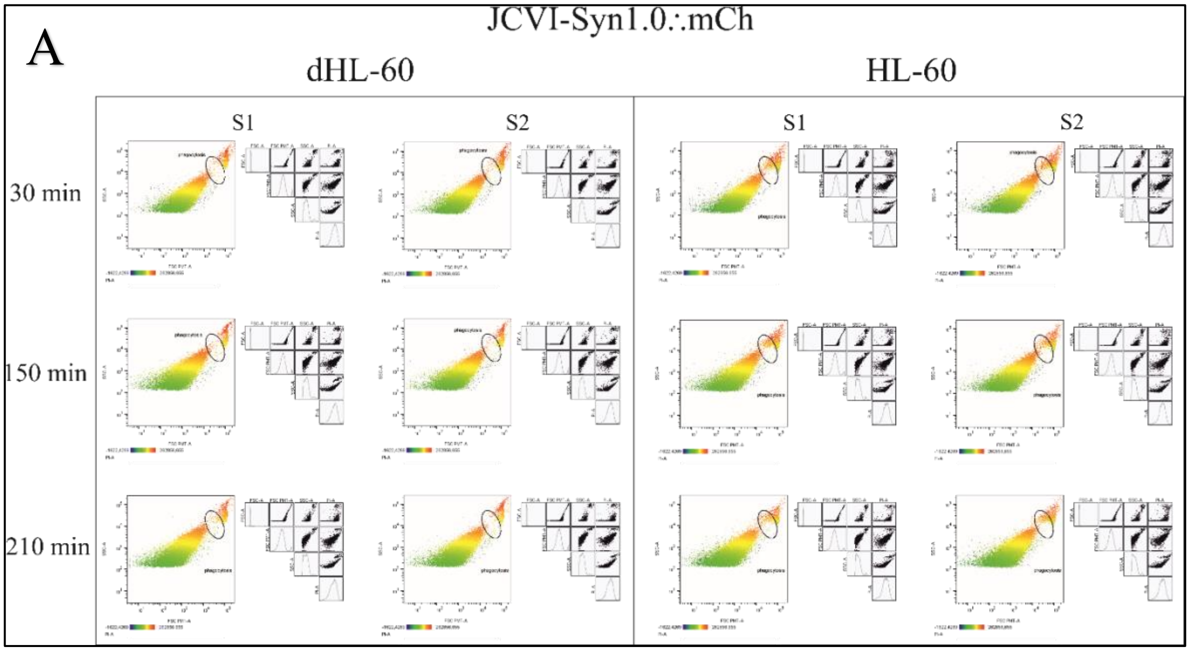


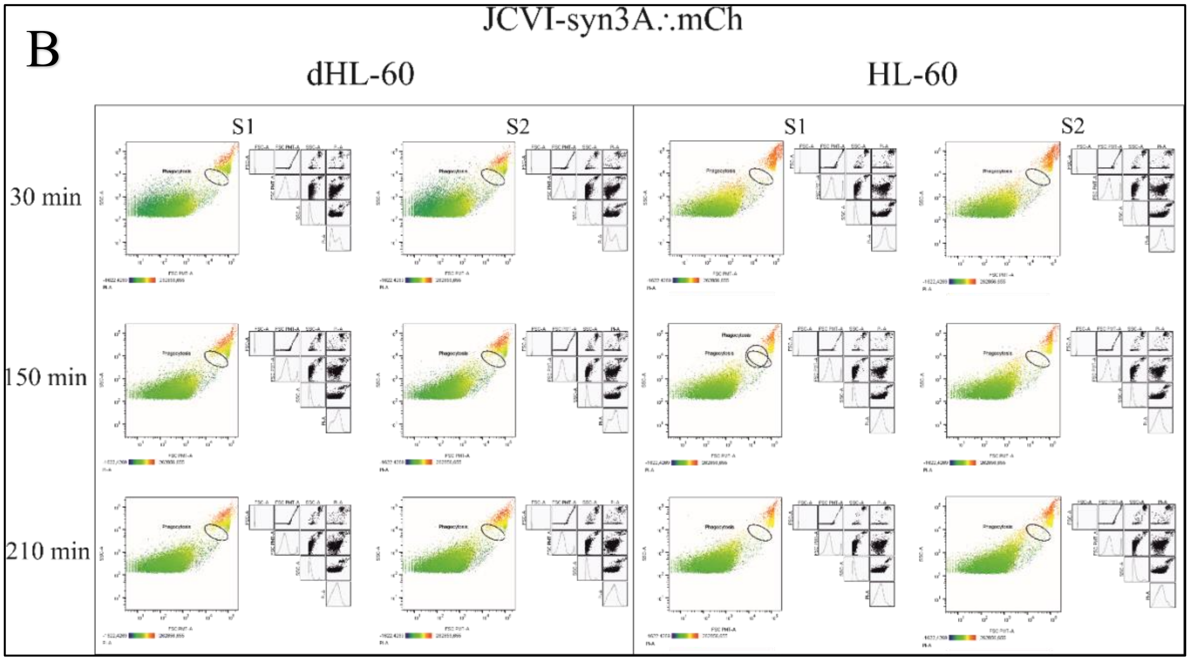

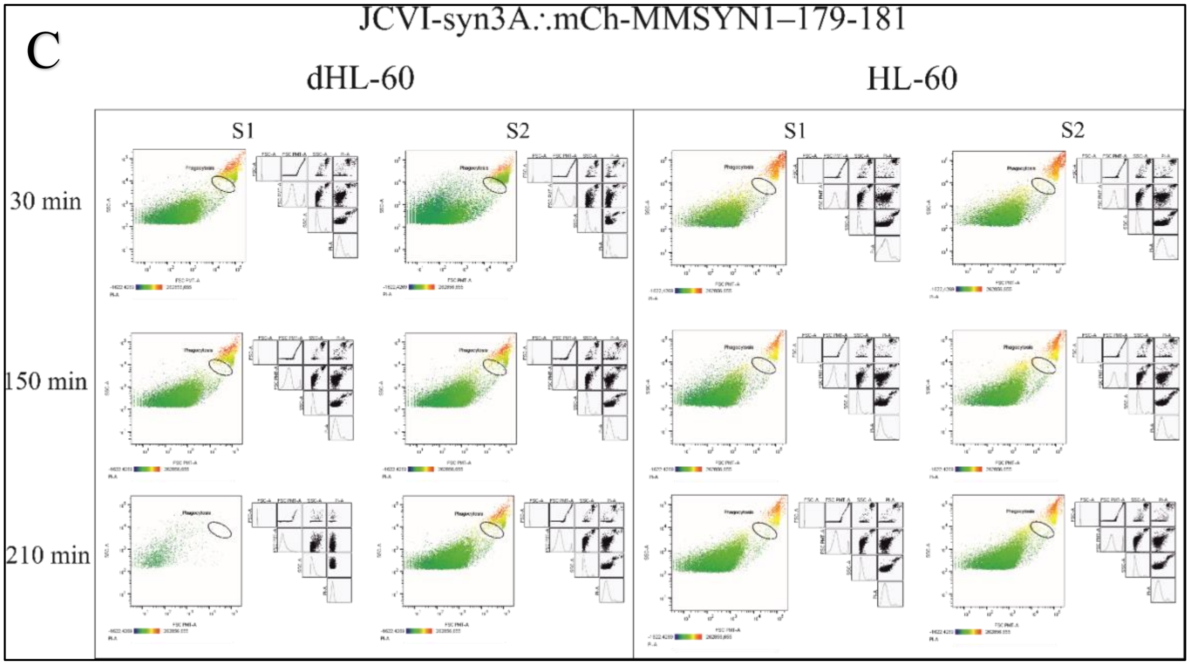
C
JCVI-syn3A∴mCh-MMSYN1–179-181
dHL-60
HL-60
S1
S2
S1
S2
30 min
150 min
210 min

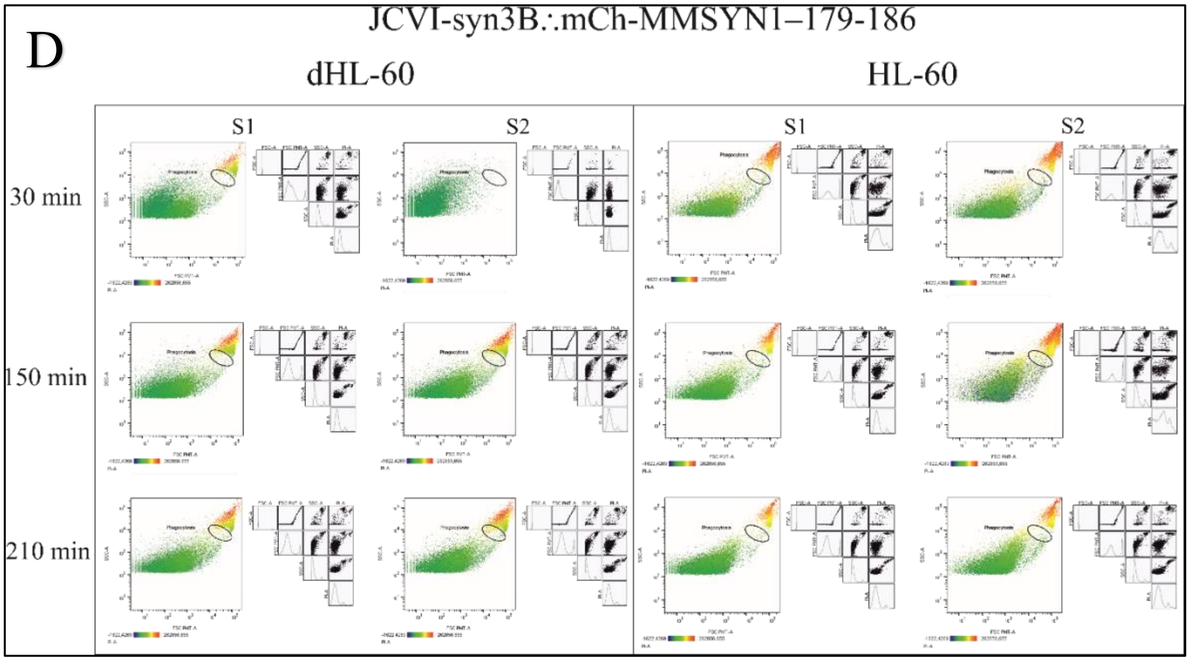
D
JCVI-syn3B∴mCh-MMSYN1–179-186
dHL-60
HL-60
S1
S2
S1
S2
30 min
150 min
210 min

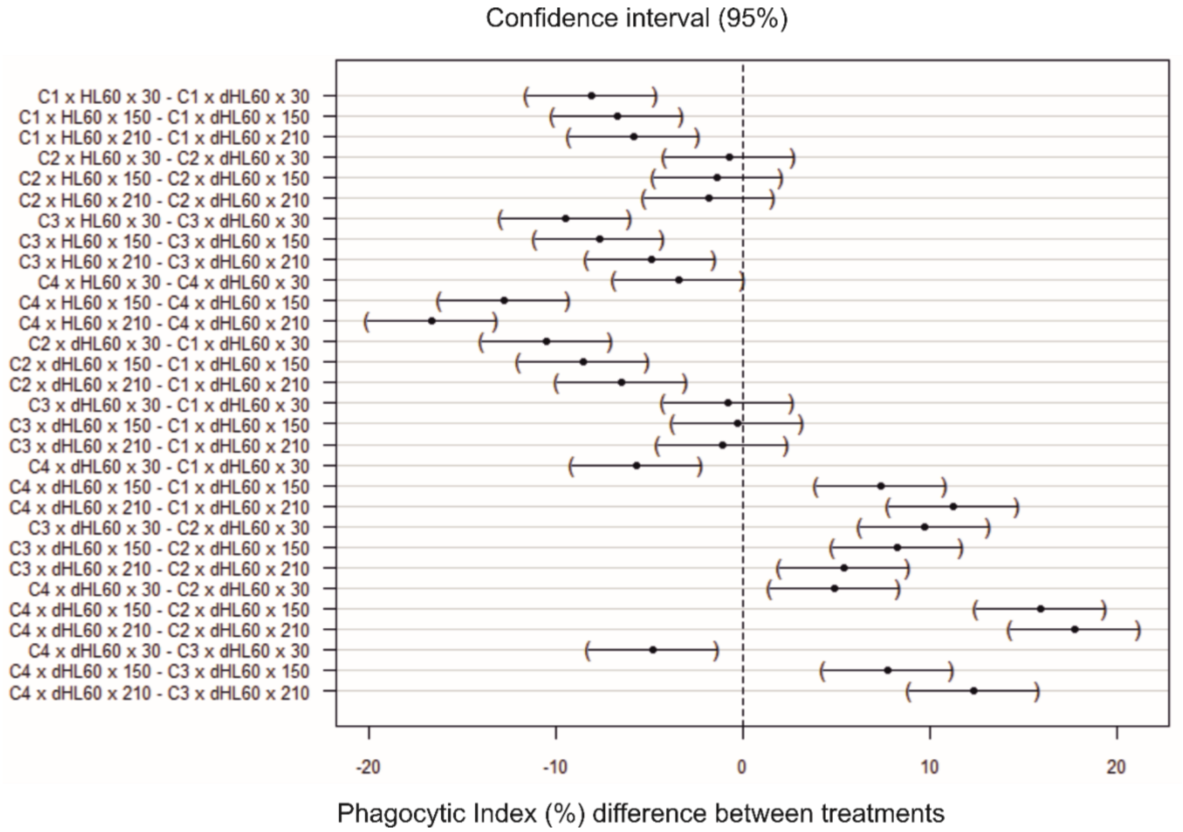


**Figure S7 – Phagocytic Index (%) difference between treatments with a confidence interval of 95%,** *related to Figure 6E*. C1, JCVI-syn1::mCh; C2, JCVI-syn3A::mCh; C3, JCVI-syn3B::mCh-MMSYN1-0179-0181 and C4, JCVI-syn3B::mCh-MMSYN1-0179-0186.
